# Experimental evolution of insect resistance to two pesticide classes reveals mechanistic diversity and context-dependent fitness costs

**DOI:** 10.1101/2021.09.03.458899

**Authors:** Stephanie S.L. Birnbaum, Nora K.E. Schulz, Destane S. Garrett, Ann T. Tate

## Abstract

Does rapid adaptation to stressors evolve through similar underlying mechanisms among diverse populations, or are there many roads to a similar phenotype? The experimental evolution of pesticide resistance in insects provides a powerful model to study the diverse evolutionary signatures of adaptation and their associated costs. Here, we selected for resistance to two pesticides (organophosphates and pyrethroids) in six field-derived populations of the red flour beetle (*Tribolium castaneum*). After several generations of selection, we performed transcriptomic analyses and measured survival, development, and fecundity in the presence and absence of pesticides to detect fitness costs of resistance evolution. All pesticide-selected populations exhibited significantly improved survival after pesticide exposure without substantial fitness costs, compared to control populations. Populations that evolved to resist organophosphates had distinct gene expression in the presence and absence of organophosphates, supporting different detoxification mechanisms and cuticular modifications among populations. In contrast, pyrethroid resistant populations demonstrated common differential expression of cytochrome P450 transcripts. Furthermore, some populations evolved similar mechanisms against both pesticides while others showed little overlap in their evolved responses, suggesting variation in potential cross-resistance phenotypes. Overall, between populations, we observed both parallel and divergent patterns in gene expression associated with acquired pesticide resistance, without ubiquitous fitness costs.

## Introduction

Understanding the molecular bases of adaptation remains a central theme in evolutionary biology. Adaptive evolution arises from standing genetic variation or *de novo* mutations that appear during selection, and patterns of evolution can vary depending on the selection environment (Barrick and Lenski, 2013). In a given set of genetically diverse populations, will exposure to a similar environment or selective pressure lead to adaptive convergence or parallelism at the genetic level, or is it more likely that each population will evolve unique genetic solutions to adapt to the environment? The formulation of quantitative theories for the evolution of adaptation depends on empirical answers to this question (Bailey and Bataillon, 2016; Barghi et al., 2020), but the adaptive significance of genetic variation in the wild can be difficult to disentangle from random noise. Experimental evolution provides a controlled approach to investigate how genetically diverse organisms adapt to the same or different selective environments. When coupled with genomic or transcriptomic analyses, researchers can monitor the course of molecular evolution in real time and on a genome-wide scale (Kawecki et al., 2012). These experiments have the power to yield important insights into the range of possible molecular mechanisms that organisms employ to improve fitness in novel environments (Barghi et al., 2020; Long et al., 2015).

One of the most ubiquitous examples of organismal adaptation to novel, stressful environments is pesticide resistance in insects. Agricultural scientists have observed insect resistance to every class of synthetic pesticide (Clark and Yamaguchi, 2002), highlighting resistance as a primary threat to sustainability of insect control. Commonly, pesticide resistance has been associated with fitness costs, but the ubiquity of these costs is under debate (ffrench-Constant and Bass, 2017; Freeman et al., 2021), partly because distinct mechanisms of resistance have variable effects on key life history fitness metrics (Assogba et al., 2015; Bajda et al., 2018; Saingamsook et al., 2019; Smith et al., 2021). For example, while target-site mutations and cytochrome P450-mediated resistance mechanisms confer similar levels of pyrethroid resistance in *Aedes aegypti*, only P450-mediated resistance was associated with fitness costs (Smith et al., 2021).

Because many synthetic pesticides act as xenobiotic stressors that impact diverse organismal functions, mechanisms that provide resistance can vary and include target-site insensitivity, metabolic detoxification, barriers to penetration, and behavioral avoidance (Boyer et al., 2012). Studies identifying resistance mechanisms, which generally rely on comparisons of field-collected susceptible and resistant populations, suggest that mechanisms of evolved resistance are hard to predict *a priori*, and may depend on standing genetic variation within a population or the shape of the fitness landscape (Adelman et al., 2011; Fang et al., 2019; Reyes et al., 2015). Resistance to organophosphates, which inhibit acetylcholinesterase (AChE) to overexcite cholinergic synapses (Casida and Durkin, 2013), and pyrethroids, which disrupt voltage-gated sodium channel function (Davies et al., 2007), provide a particularly good model for this diversity. Target-site mutations and AChE gene duplications have been described for several organophosphate-resistant insects (Siegfried and Scharf, 2001), while pyrethroid resistance has been associated with mutations in the voltage-gated sodium channel (Soderlund, 2011). Target site mutations are rarely the whole story, however, as changes to the expression of existing genes also play a substantial role in resistance evolution. Organophosphates and pyrethroids are mainly detoxified through oxidation and hydrolysis, and resistance associated with differential expression of diverse canonical detoxification genes has been described for both pesticide types (Boyer et al., 2012; Meinke et al., 2021; Oakeshott et al., 2013; Pavlidi et al., 2018; Sogorb and Vilanova, 2002). Increased resistance to penetration through changes in cuticle and serine endopeptidase gene expression have also been implicated in organophosphate and pyrethroid resistance (David et al., 2014; Lilly et al., 2016; Tandonnet et al., 2020; Zimmer et al., 2017).

Environmental heterogeneity and unknown population genetic histories can complicate interpretation of resistance mechanisms in field-collected populations and frustrate accurate estimations of relative fitness or adaptation (Feyereisen et al., 2015; Pélissié et al., 2018). Laboratory selection experiments overcome these issues by allowing for the evolution of resistance under controlled environments using distinct genetic backgrounds, yet to our knowledge, few insecticide selection studies employ the use of multiple diverse ancestral insect populations to investigate molecular resistance mechanisms (Saavedra-Rodriguez et al., 2014, 2012; Viana-Medeiros et al., 2018). In a landscape with distinct populations, are all populations capable of evolving resistance to diverse pesticides? Does selective pressure from pesticides tend to favor similar resistance solutions early in the evolutionary response, or are evolutionary trajectories likely to be diverse and population-specific? Does resistance come at a cost for all populations? By evolving multiple ancestral populations under different pesticide regimes, we can investigate the patterns of parallelism or divergence of adaptations towards the same or different stressors.

In the present study, we used an experimental evolution approach to select separately for resistance to organophosphates and pyrethroids in six distinct field-derived populations of the red flour beetle, *Tribolium castaneum*. This species is a worldwide pest of stored grains and has evolved field resistance to all classes of pesticides used for its control (David W. Hagstrum and Phillips, 2017). The *T. castaneum* populations used here have previously demonstrated constitutive variation in development time, immune gene expression, and resistance to *Bacillus thuringiensis* (Jent et al., 2019; Tate et al., 2021). Thus, we expected substantive standing genetic variation between populations and the possibility for populations to evolve pesticide resistance via diverse mechanisms. The specific goals of our study were to determine if pesticide resistance evolved in all populations, what the costs of resistance were, if the same resistance mechanisms evolve in different populations, and if these resistance mechanisms overlap. For each ancestral *T. castaneum* population, we exposed subpopulations to a control treatment or LC50 doses of pesticides (organophosphates [OP] or pyrethroids [Pyr]) (Fig. 1). After at least eight generations of selection, we compared survival, development time, and fecundity for control and evolved populations in the presence and absence of pesticides. Because we wanted to obtain a full picture of the physiological changes associated with pesticide resistance, we then used RNAseq analyses to identify gene expression changes associated with evolved resistance and pesticide exposure in three representative populations. Results from this work provide important evidence for the diverse trajectories organisms take in adapting to novel environmental stressors.

**Figure 1.**
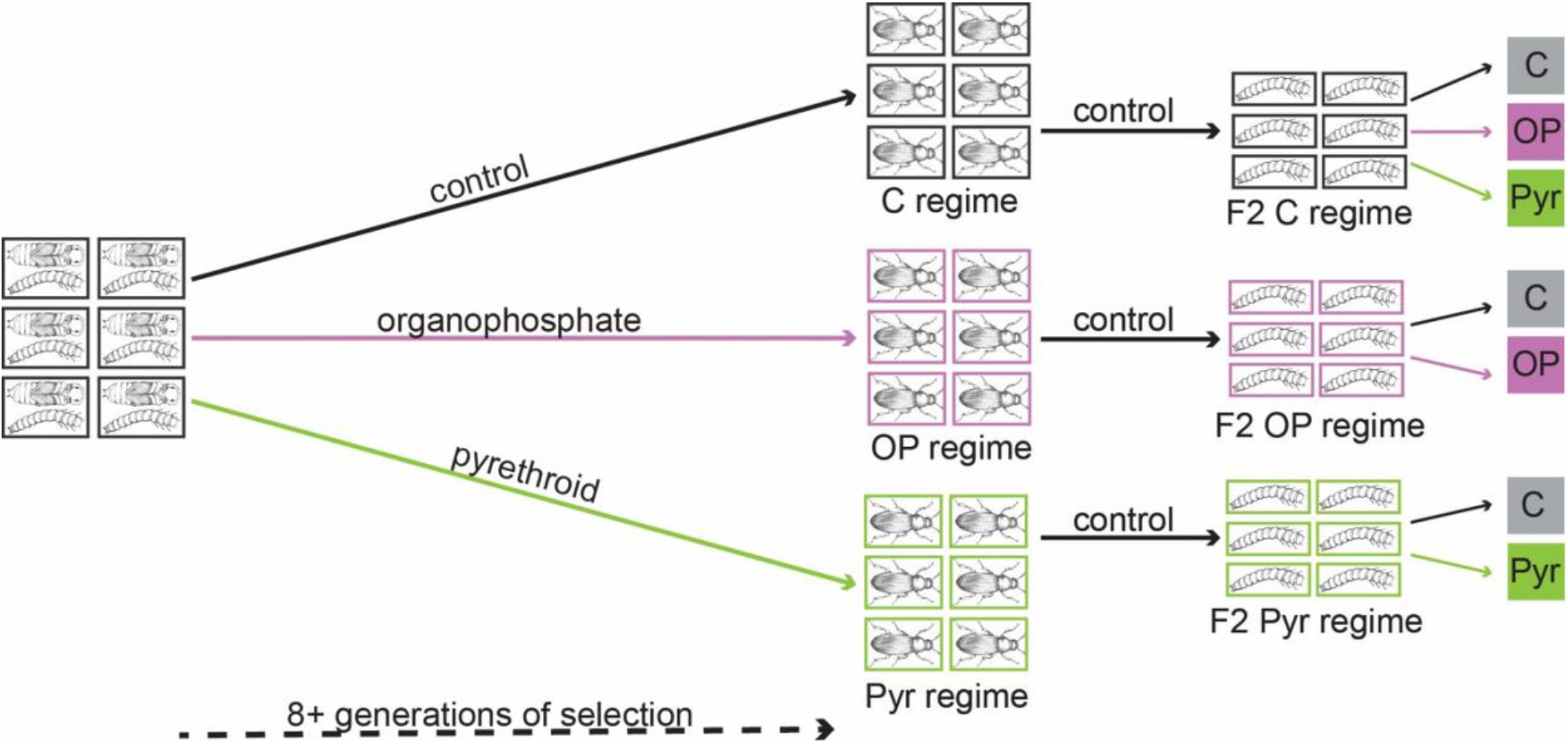
Outline of experimental design. For each of six field-collected *T. castaneum* populations, ancestral larvae and pupae were passaged for eight generations on control (C), LC50 organophosphate (OP), or LC50 pyrethroid (Pyr) selection regimes. Adults from generation 8 (or 16 for OP fecundity experiment) were allowed to produce offspring under pesticide-free (control) conditions to mitigate parental effects, and larvae at the subsequent generation (F2) were exposed to control or pesticide treatments. For the main experiment, survival and development were recorded for 21 days post-exposure for all populations. Samples for RNAseq were collected after two days of exposure for three representative populations.

## Results

### Effects of Pesticide Selection and Exposure on Survival and Development

To select for pesticide resistance, we exposed six *T. castaneum* populations (Adairville, Coffee, Dorris, RR, Snavely, and WF Ware) collected from the southeast USA (Jent et al., 2019) to either control, organophosphate (OP), or pyrethroid (Pyr) diets (Fig. 1). After eight generations of selection on either control or pesticide diets, we relaxed selection for two generations, and then exposed F2 control and pesticide-selected populations to pesticides, measured associated mortality, and analyzed the effect of selection regime on survival using coxph or parametric survival analyses. Initially, control populations varied in their susceptibility to the same dose of pesticides (Suppl. Fig. 1), with RR and Snavely populations being most susceptible to both pesticides. After eight generations of OP or Pyr selection, all selected populations exhibited significantly increased survival compared to control populations when exposed to pesticides, but varied in their relative mortality rates (Fig. 2 (Survival ∼ Regime); Suppl. Table 1). Pesticide-selected Snavely and RR populations achieved the greatest reduction in susceptibility against a given dose of both pesticides when compared to their respective control populations (Suppl. Fig. 1).

**Figure 2.**
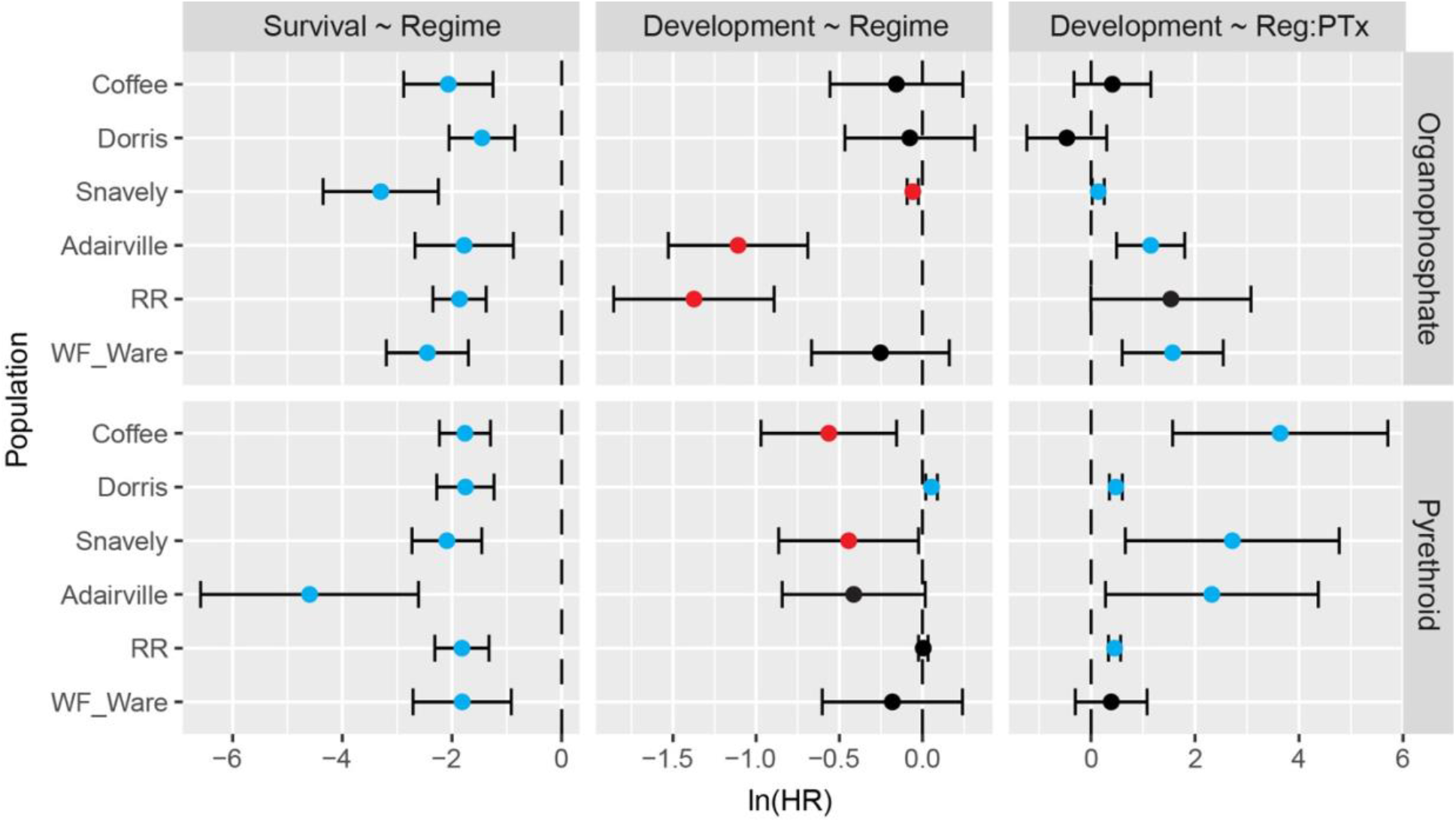
All pesticide-selected populations are more resistant to pesticides, but selection has variable costs to development. Normalized hazard ratios [ln(HR)] and 95% CI associated with survival and development models are shown (Suppl. Tables 1, 2). An HR <0 for survival indicates lower mortality, while an HR <0 for development indicates slower development. Black points are nonsignificant, blue points indicate a positive significant effect, red points indicate a negative significant effect. Survival ∼ Regime: the effect of selection regime (pesticide-selected compared to control) on survival of pesticide treated individuals; Development ∼ Regime: the effect of selection regime (pesticide-selected compared to control) on development (time to eclosion); Development ∼ Reg:PTx: the interaction effect of regime (pesticide-selected compared to control) and pesticide treatment (pesticide compared to control) on development.

In addition to negatively impacting survival, exposure to pesticides can also have deleterious impacts on development. Correspondingly, the spread of pesticide resistance through populations is influenced by the ability of resistant populations to successfully develop to reproductive age under pesticide conditions. We used coxph or parametric survival analyses for each population separately to analyze the individual and interaction effects of pesticide treatment and selection on adult development time (Day ∼ Treatment*Regime). Both OP and Pyr treatments negatively impacted development time for all control populations (Suppl. Fig. 2; Suppl. Table 2). The effects of pyrethroid exposure on development to adulthood were especially pronounced in control-regime populations, where no adults developed in any population except WF Ware (Suppl. Fig. 2B). Pyr selection mitigated the cost of pesticide exposure on development for most populations, as indicated by significant interaction effects (Fig. 2 (Development ∼ Reg:PTx); Suppl. Table 2). In particular, most Pyr-regime populations exposed to pesticides had similar development times as when reared on control diet (Suppl. Fig. 2B) indicating highly effective resistance mechanisms. The interaction effect for OP-regime and exposure was also significant for three populations (Snavely, Adairville, WF Ware), such that exposure to pesticides was less detrimental on development for OP-regime populations compared to control-regime populations (Fig. 2 (Development ∼ Reg:PTx); Suppl. Fig. 2A; Suppl. Table 2). Overall, however, pesticide exposure still delayed development for most OP-regime populations compared to development on control diets (Suppl. Fig. 2A).

To determine whether the evolution of resistance imposes a cost to development, we measured development time among selection regimes in each population in the absence of pesticide exposure. When considering the main effect of selection regime, three OP-selected populations exhibited significantly slower development compared to control populations (Snavely, Adairville, RR; Fig. 2 (Development ∼ Regime); Suppl. Fig. 2A; Suppl. Table 2). Thus, in the absence of pesticide exposure it took one (Snavely) or two days longer (Adairville and RR) to develop to adults in OP-selected relative to their control populations, which averaged around 13 days from the start of the experiment. Pyrethroid selection had mixed effects on development time, as two populations (Coffee, Snavely) exhibited significantly slower development times, while another population (Dorris) developed more quickly. However, the effect sizes were less than one day in each direction (Fig. 2 (Development ∼ Regime); Suppl. Fig. 2B; Suppl. Table 2). Thus, the evolution of organophosphate resistance was associated with more consistent costs to development time than pyrethroid resistance.

### Effects of Organophosphate Selection and Exposure on Fecundity

In a second experiment, we measured the effects of OP exposure and potential costs of OP selection on fecundity after sixteen generations of selection. Here, control and OP-selected populations were separately exposed to their own LC10 pesticide doses to minimize survivorship biases. Maternal, but not paternal, pupal weight significantly predicted fecundity when measured as live offspring (GLMM Fecundity ∼ Female weight +Male weight + (1|Population); Suppl. Fig. 3; Suppl. Table 3). This significant positive correlation was only observed in the OP exposed group (Suppl. Fig. 3A; Suppl. Table 3). Across all populations, neither OP exposure nor OP selection regime significantly predicted offspring production in the observed reproduction period. Thus, OP selection did not result in a cost to fecundity. However, there was a significant interaction of the two factors leading to higher fecundity in OP-selected individuals exposed to OP (GLMM Fecundity ∼ Regime*Tx + Female weight + (1| Population); Fig. 3; Suppl. Table 4) which could have important implications for resistant population dynamics and the spread of resistance.

**Figure 3.**
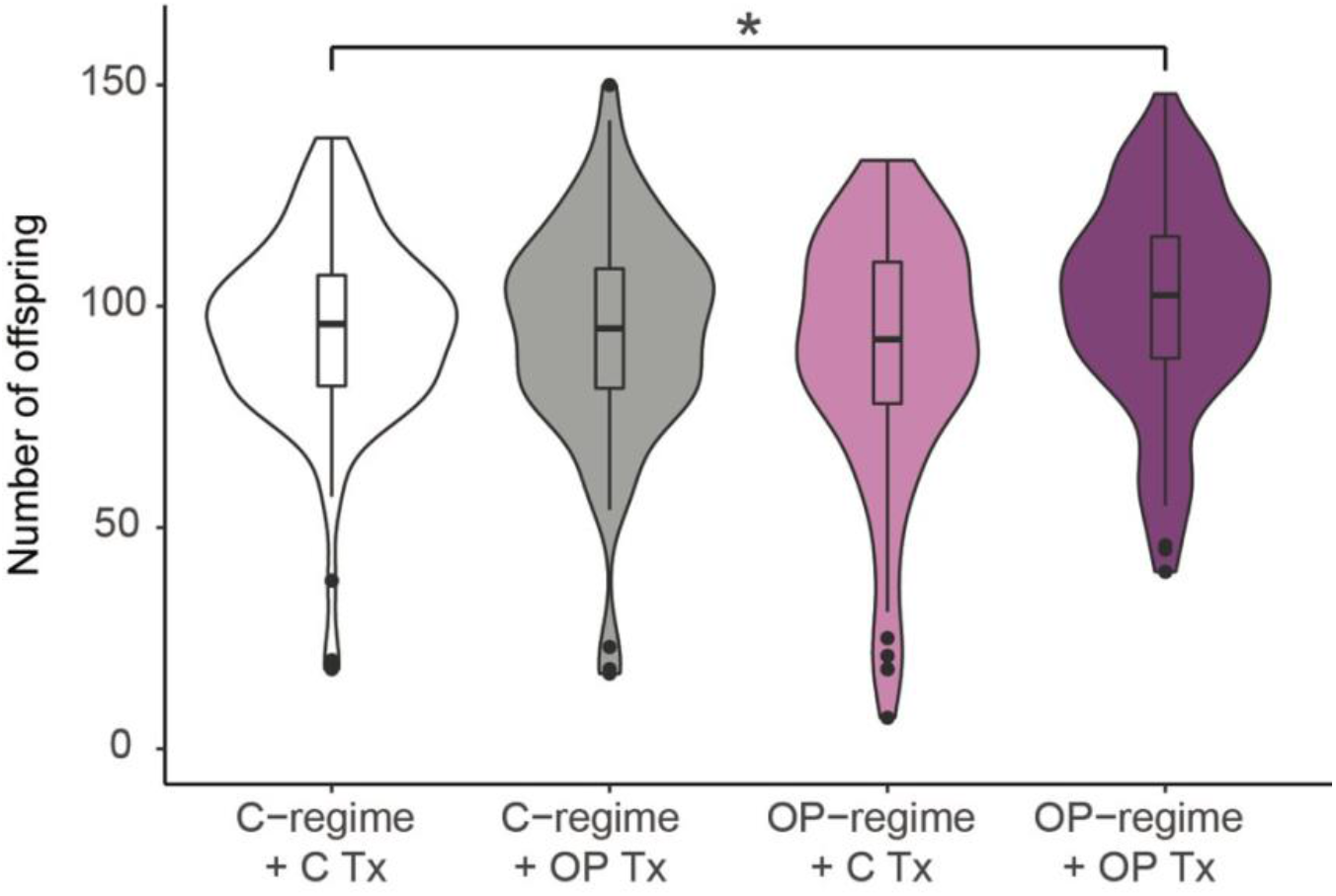
OP selection does not result in a cost to fecundity, but OP selection in combination with OP exposure during the larval stage significantly increases fecundity. The number of offspring produced from control and OP selected populations exposed to control and OP diets are shown (n ≈ 20 mating pairs per group for six populations). A significant difference between groups, as determined by GLMM analyses, is denoted by an asterisk (p < 0.05).

### Constitutive Gene Expression Differences Between Populations

We selected Coffee, Dorris, and Snavely populations to analyze the main and interactive effects of pesticide selection regime and exposure on gene expression using RNAseq analyses (sample library read counts and alignment statistics are provided in Suppl. Table 5). We collected samples for RNAseq two days after treatment exposure, before the main onset of mortality (around day four). We first analyzed constitutive differences in gene expression between populations by comparing control populations under control conditions. Coffee had the most divergent constitutive gene expression (Coffee-Dorris: 332 DE genes, Coffee-Snavely: 241 DE genes), while Dorris and Snavely had more similar constitutive gene expression (80 DE genes) (Suppl. Table 6). These results are supported by a principal component analysis (PCA) plot of control samples, wherein two Coffee replicates are separated along a component that explains 68% of the variation between unexposed control-regime samples (Suppl. Fig. 4A). PCA plots that include all regime and pesticide treatment samples demonstrated a similar pattern (Suppl. Figs. 4B, 4C), and overall, replicates from populations evolved to resist pesticides grouped closely to unexposed samples, even after pesticide exposure.

### Effects of Pesticide Exposure on Gene Expression

We predicted that populations would respond similarly to pesticide exposure, in that they would all exhibit broad differential regulation of physiological processes and differentially express a similar suite of metabolic detoxification genes relative to their unexposed controls. The results suggest that pesticide treatment did have a substantive effect on overall gene expression regardless of selection regime (Figs. 4A, D; Suppl. Figs. 5-7). Relative to unexposed beetles, beetles from Coffee, Dorris, and Snavely significantly differentially expressed 471, 412, 1241 genes in response to OP treatment and 255, 729, 435 genes in response to Pyr treatment, respectively (p < 0.05 after FDR correction; Suppl. Fig. 5).

**Figure 4.**
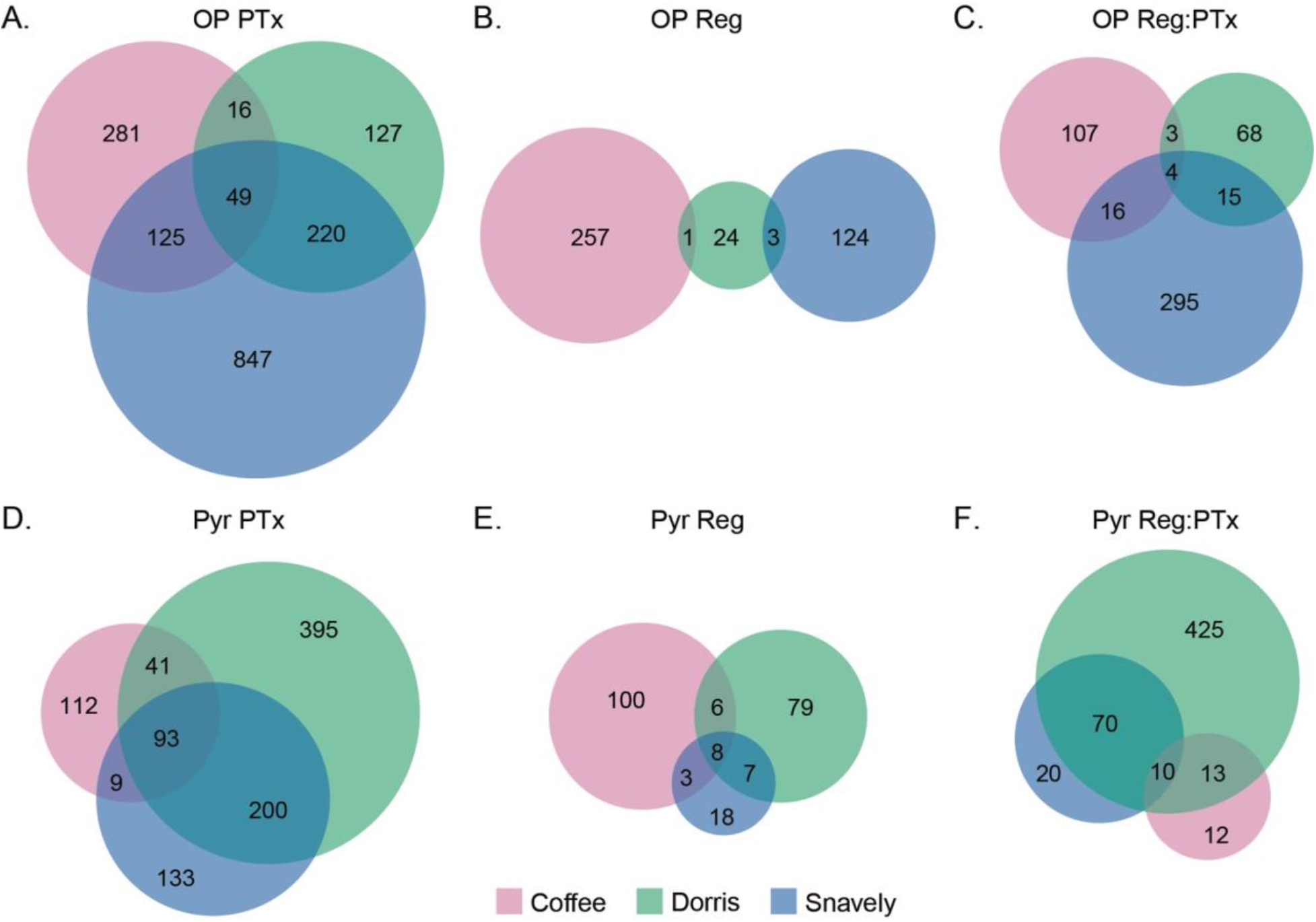
The overlap of differentially expressed genes between populations for each DE factor varies between pesticide types. Venn diagrams show the number of significantly differentially expressed genes (padj < 0.05 after FDR correction) for each population with: **A.** OP PTx: organophosphate exposure treatment, **B.** OP Reg: organophosphate selection regime, **C.** OP Reg:PTx: interaction between organophosphate selection and exposure, **D.** Pyr PTx: pyrethroid exposure treatment, **E.** Pyr Reg: pyrethroid selection regime, **F.** Pyr Reg:PTx: interaction between pyrethroid selection and exposure.

In response to OP treatment, populations commonly differentially expressed 49 genes including several canonical detoxification genes: three downregulated cytochrome P450s, one upregulated UDP-glucuronosyltransferase (UGT), and one upregulated ABC transporter (Fig. 4A; Fig. 5; Suppl. Table 6). GO enrichment analysis of genes differentially expressed in response to OP treatment revealed over 100 significantly enriched terms for each population, and the most significant molecular GO terms were “structural constituent of cuticle”, “oxidoreductase activity”, and “catalytic activity” for Coffee, Dorris, and Snavely, respectively (Fig. 6; Suppl. Table 7).

**Figure 5.**
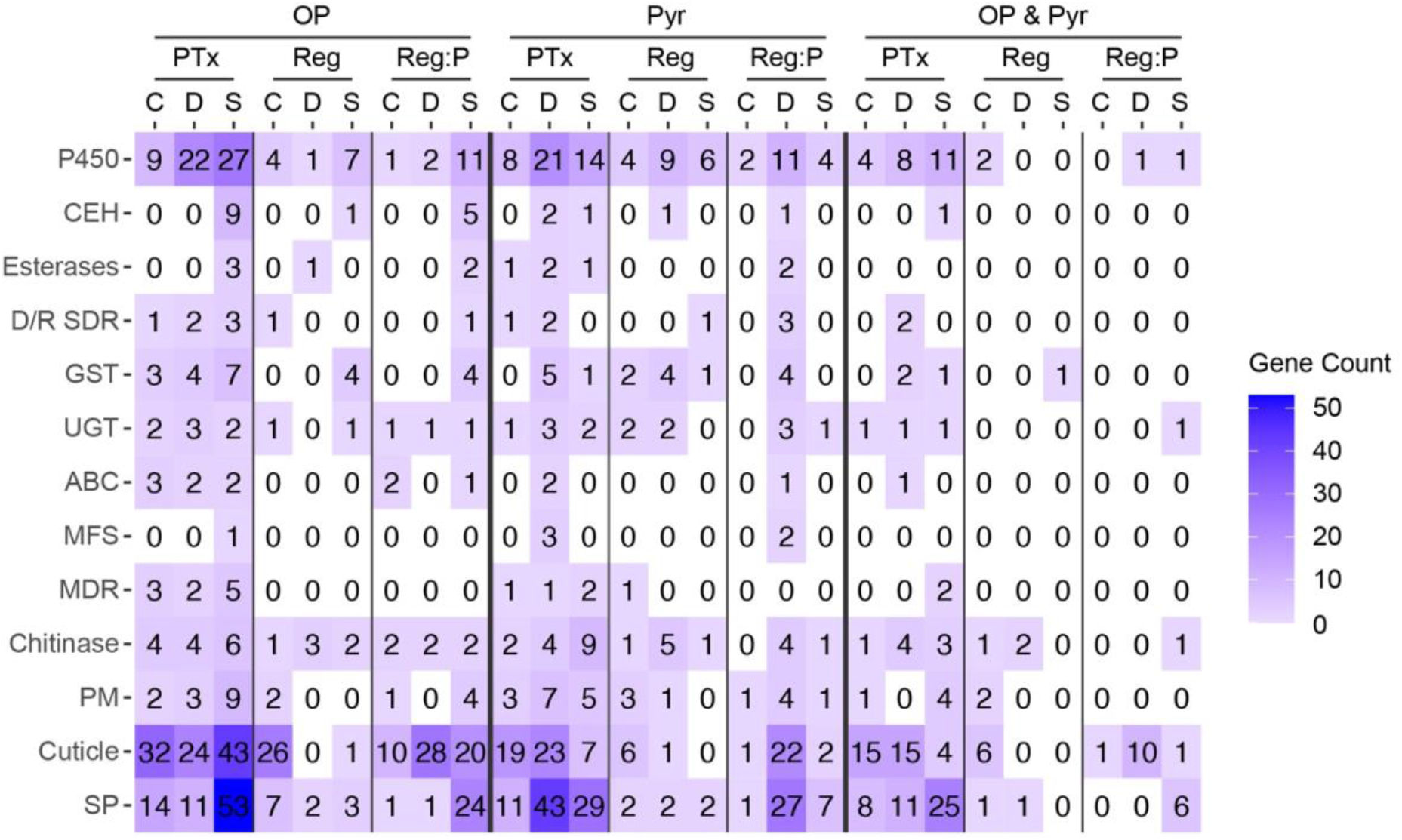
Number of genes significantly differentially expressed in selected gene groups canonically associated with pesticide resistance (padj < 0.05 after FDR correction). Cells are gradient colored to reflect the number of differentially expressed genes in each group. Abbr. and gene groups: OP = genes differentially expressed with organophosphate treatment and regime, Pyr = genes differentially expressed with pyrethroid treatment and regime, OP & Pyr = genes commonly differentially expressed among both organophosphate and pyrethroid treatments and regimes; PTx = pesticide treatment (base level is control), Reg = evolution regime (base level is control regime), Reg:P = the interaction of evolution regime (pesticide-selected compared to control) and pesticide treatment (pesticide compared to control); C = Coffee, D = Dorris, S = Snavely; P450 = cytochrome P450; CEH = carboxyl ester hydrolase, D/R SDR = dehydrogenase/ reductase SDR, MFS = MFS-type transporter, MDR = multidrug resistance protein, chitinase = chitin deacetylase & chitinase, PM = peritrophic matrix & peritrophins, SP = serine protease & serpin peptidase inhibitor.

**Figure 6.**
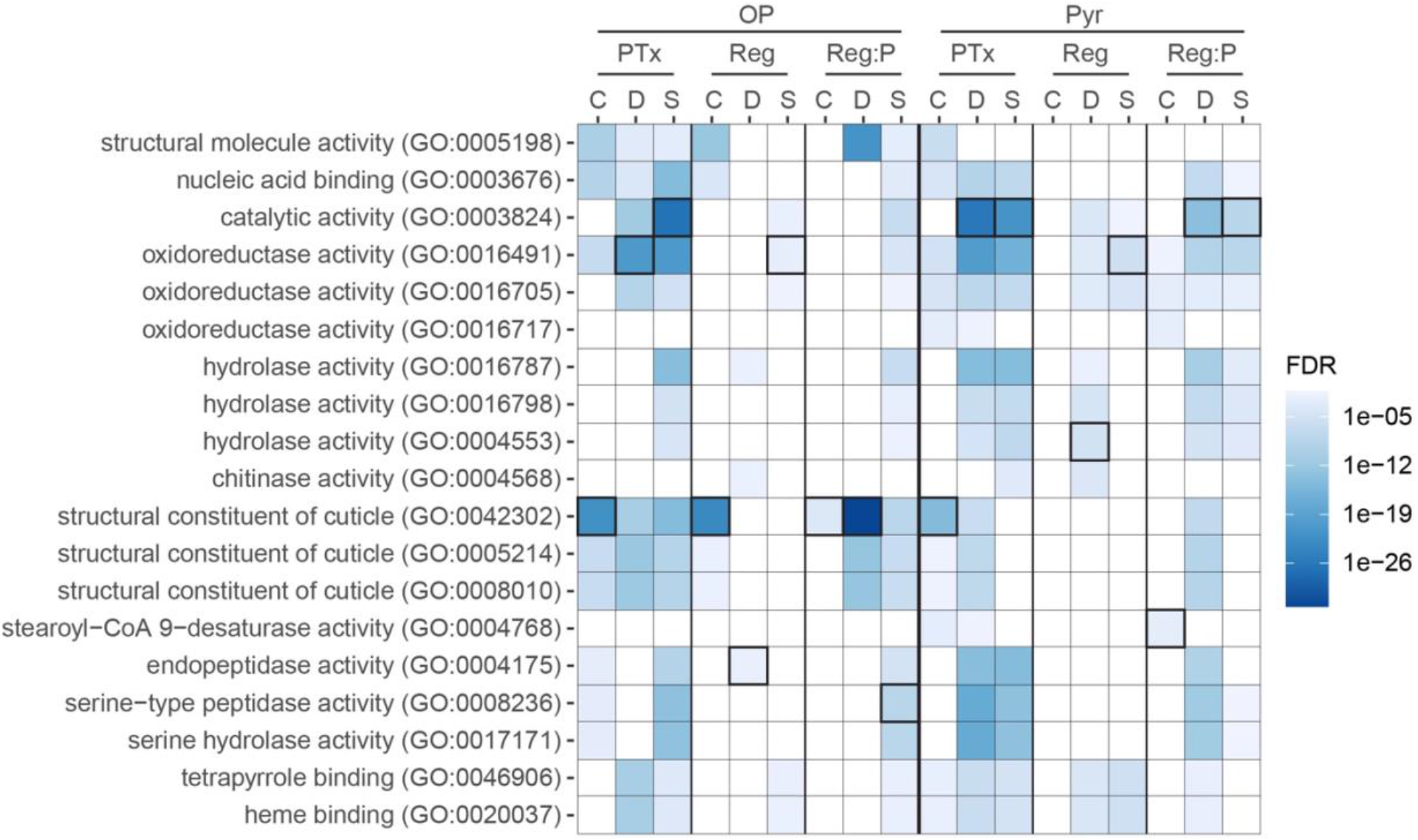
Significantly enriched molecular GO terms for each set of differentially expressed genes. The three most significantly enriched GO terms for each group were selected (the top term for each group is indicated by black outlines) and cells are shaded by false discovery rate (FDR)-adjusted p-values for all groups (non-significant terms are white). OP= organophosphate, Pyr = pyrethroid; PTx = pesticide treatment (base level is control), Reg = evolution regime (base level is control regime), Reg:P = (pesticide-selected compared to control) and pesticide treatment (pesticide compared to control); C = Coffee, D = Dorris, S = Snavely.

In response to Pyr treatment, populations commonly differentially expressed 93 genes including four P450s (two upregulated, two downregulated), one upregulated UGT, and one upregulated potassium channel subfamily K (Fig. 4D; Fig. 5; Suppl. Table 6). GO enrichment analysis of genes differentially expressed in response to Pyr treatment revealed between 87-262 significantly enriched terms for each population, and the most significantly enriched molecular GO terms were “structural constituent of cuticle” for Coffee and “catalytic activity” for Dorris and Snavely (Fig. 6; Suppl. Table 8). With a total of 410 and 343 differentially expressed genes shared between at least two of three populations in response to OP and Pyr exposure treatments, respectively, there was a high degree of similarity between populations in response to each pesticide treatment.

### Effects of Pesticide Selection on Gene Expression in the Absence of Pesticides

We predicted that selection for pesticide resistance would result in similar constitutive changes to detoxification and cuticle-related gene expression among different populations in the absence of pesticide exposure. While we did observe constitutive differential gene expression in each OP-selected population relative to its control (Coffee: 258; Dorris: 28; Snavely: 127 DE genes; Suppl. Fig. 5), no genes were commonly differentially expressed between all three populations in response to organophosphate selection, and very few were shared even among two populations (Fig. 4B; Suppl. Fig. 6; Suppl. Table 6). In contrast to the divergent responses among OP-selected populations, Pyr-selected populations exhibited a greater degree of similarity. Eight differentially-expressed genes were shared among all three populations in response to pyrethroid selection, and sixteen total genes were shared pairwise between populations, even though the overall numbers of differentially expressed genes were generally lower in pyrethroid-selected populations relative to controls (Coffee:117; Dorris: 100; Snavely: 36 DE genes; Fig. 4E; Suppl. Fig. 7; Suppl. Table 6).

Based on previous studies investigating molecular mechanisms of organophosphate resistance, we expected common differential expression of detoxification genes including cytochrome P450s, carboxyl ester hydrolases (CEHs), esterases, and glutathione S-transferases (GSTs) among OP-regime populations (Boyer et al., 2012; Meinke et al., 2021; Oakeshott et al., 2013; Pavlidi et al., 2018; Sogorb and Vilanova, 2002; Tandonnet et al., 2020). However, after organophosphate selection, each population tended to differentially express different P450, GST, and UGT genes canonically associated with detoxification (Figs. 5, 7A). Changes in cuticular metabolism can also aid in pesticide resistance (Balabanidou et al., 2018; Tandonnet et al., 2020), and Coffee, in particular, constitutively upregulated several cuticular proteins in response to organophosphate selection (Figs. 5, 8A). Interestingly, most of the cuticular genes differentially expressed after organophosphate selection in Coffee were also constitutively differentially expressed in control-regime Coffee relative to the other two populations (although generally at lower levels than the selected line). The most significantly enriched molecular GO terms reveal differences between populations in gene expression changes associated with organophosphate selection: “structural constituent of cuticle”, “endopeptidase activity”, and “oxidoreductase activity” were the top GO terms for OP-regime Coffee, Dorris, and Snavely, respectively (Fig. 6; Suppl. Table 7).

**Figure 7.**
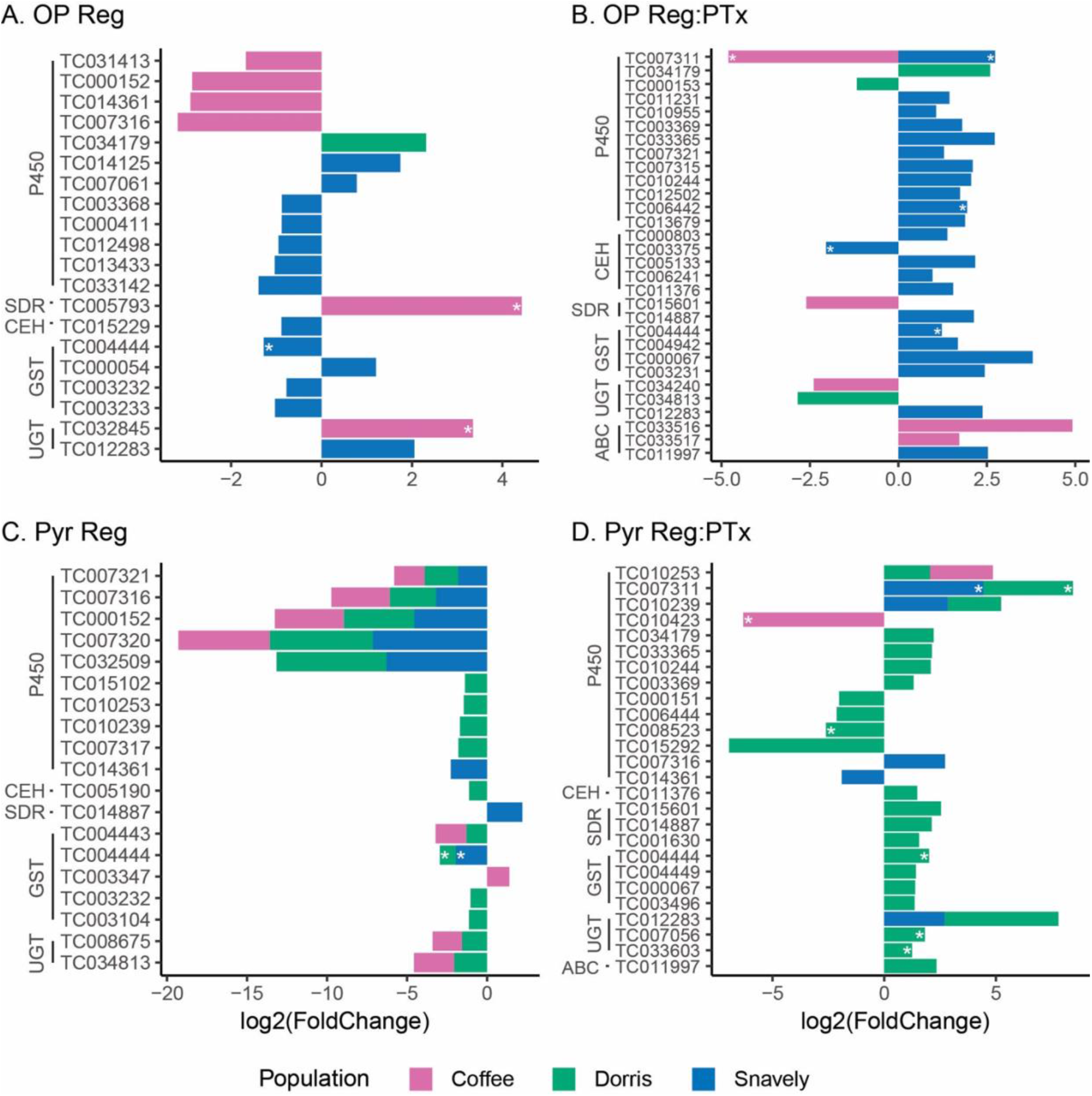
Canonical detoxification genes significantly differentially expressed after pesticide selection and the interaction between selection and pesticide exposure (padj < 0.05 after FDR correction). Stacked bar graphs show the log_2_ fold change values for: **A.** Genes DE between control and organophosphate-selected populations, **B.** Genes DE between control and organophosphate-selected populations when exposed to organophosphate treatments, **C.** Genes DE between control and pyrethroid-selected populations, **D.** Genes DE between control and pyrethroid-selected populations when exposed to pyrethroid treatments. Transcripts that were constitutively DE in pairwise population comparisons (e.g. that showed significant standing variation in the absence of pesticide selection and treatment) are further denoted with asterisks. P450 = cytochrome P450s, CEH = carboxylic ester hydrolases, SDR = dehydrogenase/ reductase SDRs, GST = glutathione S-transferases, UGT = UDP-glucuronosyltransferase, ABC = ABC transporters.

**Figure 8.**
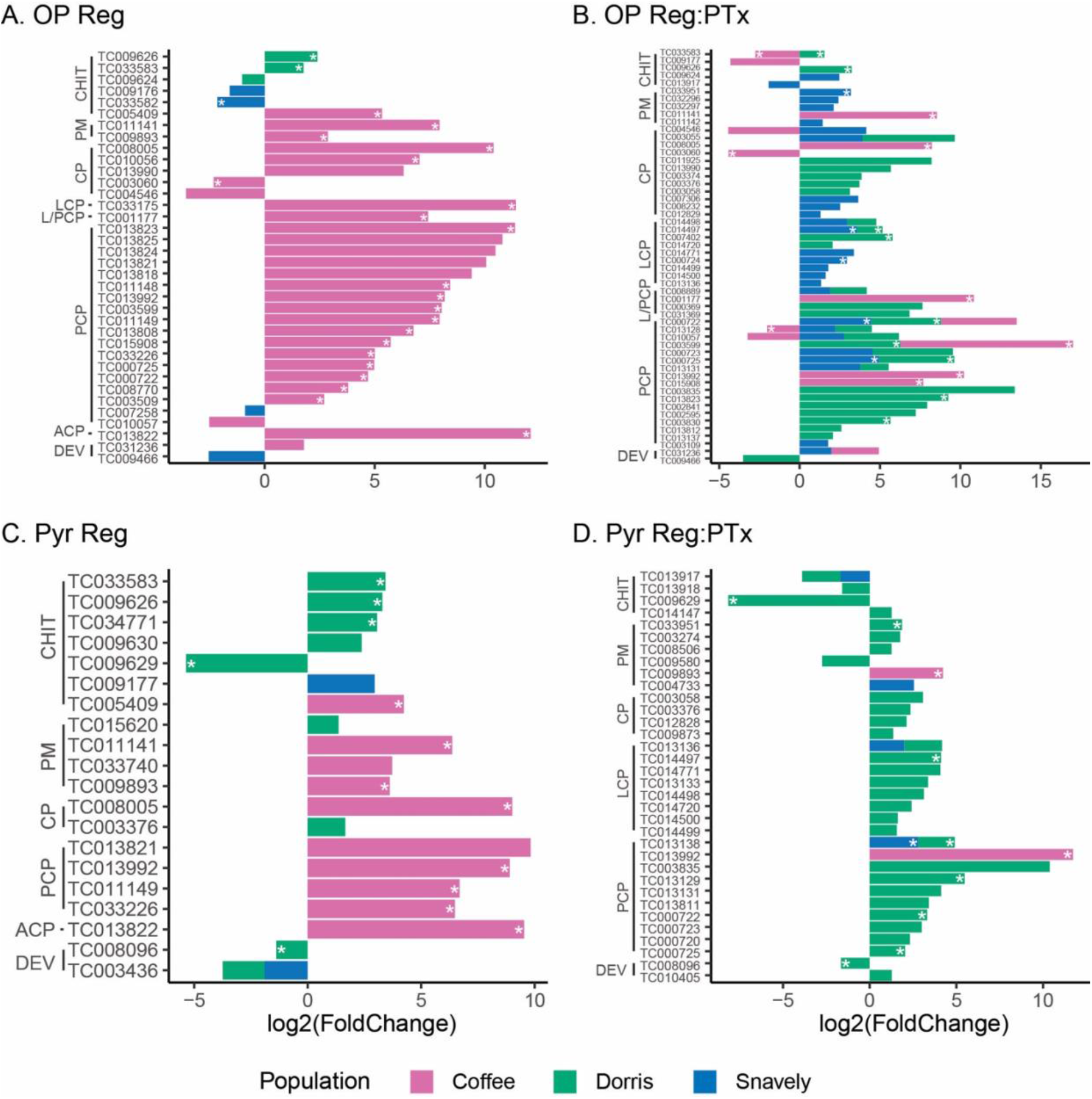
Cuticular and development genes significantly differentially expressed with pesticide selection and the interaction between selection and exposure (padj < 0.05 after FDR correction). Stacked bar graphs show the log_2_ fold change values for: **A.** Genes DE between control and organophosphate-selected populations, **B.** Genes DE between control and organophosphate-selected populations when exposed to organophosphate treatments, **C.** Genes DE between control and pyrethroid-selected populations, **D.** Genes DE between control and pyrethroid-selected populations when exposed to pyrethroid treatments. Transcripts that were constitutively DE in pairwise population comparisons (e.g. that showed significant standing variation in the absence of pesticide selection and treatment) are further denoted with asterisks. CHIT = chitin related genes, PM = peritrophic matrix related genes, CP = cuticular proteins, LCP = larval cuticular proteins, L/PCP = larval/ pupal cuticular proteins, PCP = pupal cuticular proteins, ACP = adult cuticular proteins, DEV = core development genes.

Previous studies have highlighted the importance of P450s, esterases, GSTs, and other canonical detoxification genes underlying pyrethroid resistance (Boyer et al., 2012; David et al., 2014; Meinke et al., 2021; Zimmer et al., 2017). Indeed, we found that Pyr-regime populations commonly downregulated several canonical detoxification genes including five P450s, two GSTs, and two UGTs (Figs. 5, 7C). Pyr-regime Coffee upregulated fewer cuticular genes in response to pyrethroid selection compared to selection with organophosphates (Figs. 5, 8C). Again, many of the cuticular genes differentially expressed between control- and Pyr-regime Coffee and Dorris populations were also constitutively differentially expressed in pairwise comparisons between control-regime populations. Dorris and Snavely Pyr-regime populations were enriched for differentially expressed genes involved the molecular functions of “hydrolase activity” and “oxidoreductase activity”, respectively (Fig. 6; Suppl. Table 8); genes differentially expressed in Pyr-regime Coffee revealed no significantly enriched molecular GO terms. Overall, there were more similarities in gene expression responses among populations after pyrethroid selection compared to selection with organophosphates.

### Differential Gene Expression of Pesticide-Selected Populations when Exposed to Pesticides

Among populations, the overall concordance of genes differentially expressed between control and pesticide-selected populations after exposure to pesticides (the interaction term) can provide insight into whether the evolution of resistance arises through parallel or different mechanisms. Relative to the main effect of selection (discussed above), there was greater overlap between populations in the genes differentially expressed with respect to the interaction term, suggesting that responses induced by pesticides show a greater degree of parallelism than constitutive states among resistant populations. Again, however, the organophosphate-associated populations were more dissimilar than the pyrethroid-associated populations (Figs. 4C, F).

Beetles selected with OP exhibited large differences in the number of differentially expressed genes relative to control beetles after pesticide exposure (Coffee: 130; Dorris: 90; Snavely: 330 DE genes, Suppl. Fig. 5); only four genes were shared between all populations, and 34 genes were shared pairwise between populations (Fig. 4C, Suppl. Fig. 6). No canonical detoxification genes (Casida, 2017; Feyereisen et al., 2015; Panini et al., 2016) were commonly differentially expressed between any of the populations (Fig. 7B). Dorris differentially expressed only two P450s and one UGT, and Coffee differentially expressed one P450, one short-chain dehydrogenases/reductase (SDR), one UGT, and two ABC transporters. In contrast, Snavely generally upregulated over twenty detoxification genes including several cytochrome P450s, CEHs, and GSTs, (Figs. 5, 7B). All three populations differentially expressed a number of different cuticle-related genes, and Dorris and Snavely upregulate a greater number of cuticular genes in response to the interaction between OP selection and treatment compared to Coffee (Figs. 5, 8B). GO enrichment analyses of OP selection (regime)-by-treatment differentially expressed genes indicated the importance of molecular functions in “structural constituent of cuticle” for Coffee and Dorris and “serine-type peptidase activity” for Snavely (Fig. 6; Suppl. Table 7).

In response to pyrethroid exposure, Pyr-selected populations exhibited greater similarity in gene expression, with 93 genes shared between at least two populations (Fig. 4F; Suppl. Fig. 7). Coffee and Dorris commonly upregulated one P450 and Dorris and Snavely shared in the overexpression of two P450s and one UGT (Figs. 5, 7D). In particular, Dorris Pyr-regime populations differentially expressed several detoxification and cuticular genes in response to pyrethroid exposure (Figs. 5, 7D, 8D). GO enrichment analyses revealed the importance of “stearoyl-CoA 9-desaturase activity” for Coffee and “catalytic activity” for Dorris and Snavely among Pyr regime:treatment differentially expressed genes (Fig. 6; Suppl. Table 8). Overall, there was also greater similarity between populations in genes responding to the interaction between pyrethroid selection regime and exposure compared to genes differentially expressed with organophosphate selection and exposure.

### Overlap of Gene Expression Changes in Response to Organophosphates and Pyrethroids

We wanted to quantify the overlap between responses to organophosphates and pyrethroids, as metabolic detoxification genes implicated in interactions with both pesticides could point to the possibility of cross-resistance or facilitate evolution of resistance to multiple pesticides. Given the large number of genes differentially expressed after pesticide exposure, it is not surprising that a large numbers of differentially expressed genes were shared after treatment with OP and Pyr (Snavely: 307; Dorris: 254; Coffee: 154 genes DE in response to each pesticide exposure; Figs. 9A-C). Populations varied in the number of detoxification genes commonly differentially expressed with OP and Pyr treatments (Fig. 5). However, in response to both pesticide treatments, all three populations differentially expressed several P450s, cuticle-related, and serine protease genes, which may compose a core of non-specific detoxification responses.

**Figure 9.**
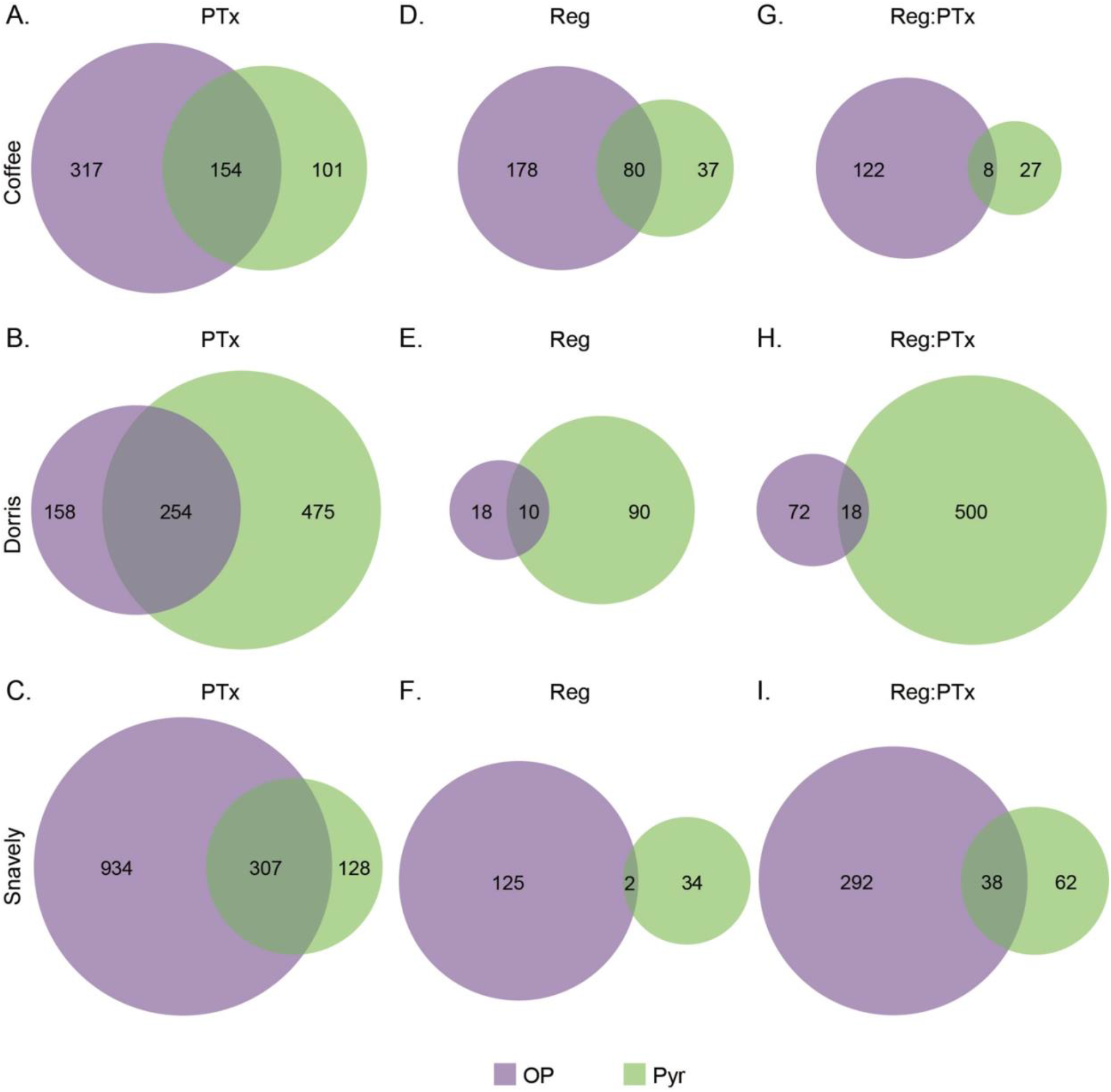
Overlap of significantly differentially expressed genes between organophosphate and pyrethroid pesticide treatments and regimes for each population (padj < 0.05 after FDR correction). Pesticide exposure: **A.** Coffee, **B.** Dorris, **C.** Snavely; selection regime: **D.** Coffee, **E.** Dorris, **F.** Snavely; interaction between pesticide selection and exposure: **G.** Coffee, **H.** Dorris, **I.** Snavely.

Genes commonly differentially expressed after selection against organophosphates and pyrethroids could point to mechanisms that enable the evolution of cross-resistance. Populations varied dramatically in the number of differentially expressed genes shared between OP and Pyr selection regimes with 80, 10, and 2 genes shared between pesticide-selected Coffee, Dorris, and Snavely populations, respectively (Figs. 9D-F). Populations were especially varied in the canonical detoxification, cuticular, and serine protease genes shared between OP- and Pyr- regimes (Fig. 5). Of the genes significantly differentially expressed with respect to the interaction between pesticide selection regime and exposure, Coffee, Dorris, and Snavely shared 8, 18, and 38 genes, respectively (Figs. 9G-I). Again, specific detoxification, cuticular, and serine protease genes played variable roles in genes commonly responding to the interactions between pesticide regime and exposure (Fig. 5). Assuming that similar gene expression patterns after pesticide exposure could predict or facilitate the evolution of cross-resistance, the potential for cross-resistance appears to vary among populations.

## Discussion

The diversity of adaptive mechanisms among populations and the evolutionary landscape of adaptation in response to a multitude of stressors remain central themes in evolutionary biology. The polygenic responses underlying pesticide resistance to diverse classes of pesticides and the degree of parallelism among resistance mechanisms in different populations present an opportunity to understand the evolutionary processes underlying the scourge of pesticide resistance. Adaptation to novel environments or stress is often associated with polygenic changes and mechanistic variation among populations (Barghi et al., 2020; Jain and Stephan, 2017). Understanding this variable genetic architecture has important implications for the long-term evolution of diverse taxa (Bullard et al., 2010; Dowle et al., 2020; Harrisson et al., 2014; Rose et al., 2018), and importantly, for the spread of pesticide resistance (ffrench-Constant, 2013; McKenzie and Batterham, 1994). Here, we selected for resistance to organophosphates and pyrethroids in several *T. castaneum* populations and compared fitness component and gene expression responses to pesticide selection and exposure. All pesticide-selected populations had increased survival compared to control populations when exposed to pesticides, but there were variable selection-associated costs to development time and fecundity among pesticides and populations. Pesticide-selected populations varied in the genes differentially expressed with respect to selection and exposure. However, there was greater similarity of gene expression responses with pyrethroid interactions compared to organophosphate gene expression responses, demonstrating that parallel changes in gene expression between populations are dependent on the stressor. Furthermore, the genes commonly differentially expressed against both pesticides varied between populations.

Past studies examining gene expression and enzymatic differences between susceptible and resistant populations have indicated the importance of a number of metabolic mechanisms underlying organophosphate and pyrethroid resistance. Organophosphate resistance has been most widely associated with acetylcholinesterase (AChE) copy number variation or target-site modifications and metabolic detoxification by carboxyl ester hydrolases (CEHs) and other esterases (Alon et al., 2008; Carletto et al., 2010; Carvalho et al., 2013; Coates et al., 2016; Haubruge et al., 2002; Kwon et al., 2010). Differential expression of cytochrome P450s, glutathione S-transferases (GSTs), short-chain dehydrogenases/reductases (SDRs), serine-endopeptidases, and cuticular genes have also been described in organophosphate resistant insect populations (Carvalho et al., 2013; Saavedra-Rodriguez et al., 2014; Tandonnet et al., 2020; Zhu et al., 2012). In particular, other studies examining population variation in organophosphate resistance mechanisms have found common differential activity of esterases (Reyes et al., 2015) and GSTs (Saavedra-Rodriguez et al., 2014). We found that populations had divergent changes in gene expression in response to organophosphate selection. Several cytochrome P450s, CEHs, SDRs, GSTs, UDP-glucuronosyltransferase (UGTs), and ABC transporters were differentially expressed in response to organophosphate selection and the interaction between selection regime and organophosphate exposure, but none of these canonical detoxification genes were commonly differentially expressed between any of the populations. Dorris and Snavely organophosphate-selected populations upregulated expression of an acetylcholinesterase-like protein when exposed to organophosphates, but Snavely alone differentially expressed CEHs in response to organophosphate selection and exposure. Dorris and Snavely, while differentially expressing different genes, appear to rely on endopeptidase and oxidoreductase activity as illustrated by the differential expression of different serine proteases and P450s. Organophosphate-resistant Coffee individuals may especially rely on cuticular modifications, as indicated by GO enrichment analyses of genes differentially expressed with respect to regime and the interaction between regime and pesticide treatment.

In addition to target-site modifications (Davies et al., 2007; Silva et al., 2014), pyrethroid resistance has also been associated with a number of canonical detoxification genes including many P450s, CEHs, SDRs, GSTs, UGTs, ABC transporters, and cuticular proteins (Adelman et al., 2011; Carletto et al., 2010; Carvalho et al., 2013; David et al., 2014; Kostaropoulos et al., 2001; Saavedra-Rodriguez et al., 2012; Yan et al., 2018; Zimmer et al., 2017). Here, we identified a variety of cytochrome P450s, CEHs, SDRs, GSTs, UGTs, and ABC transporters that were differentially expressed in response to pyrethroid selection and exposure and potentially involved in resistance mechanisms. One previous study examining the molecular bases of pyrethroid resistance among *A. aegypti* populations found that very few genes commonly responded between populations to pyrethroid selection (Saavedra-Rodriguez et al., 2012). In contrast, our results suggest that pyrethroid selection yielded a substantial number of parallel gene expression responses between populations. In congruence with past evidence for the importance of P450s in pyrethroid resistance (Scott et al., 2015; Yan et al., 2018), all three Pyr- regime populations commonly constitutively downregulated four cytochrome P450 genes (6BQ11, 6BQ7, 345A1, 6BQ10). While pesticide resistance is often associated with the overexpression of P450s, other studies have observed the downregulation of P450s in insecticide resistant populations (David et al., 2014; Tandonnet et al., 2020; Yang and Liu, 2011). Decreased expression of specific P450s might reduce the metabolism of compounds into more active or damaging forms or be involved in controlling inflammatory responses (Feyereisen, 2019; Wang et al., 2017).

Cuticular modifications can reduce the penetration of pesticides through the exoskeleton or gut and slow insecticide action within the insect (Balabanidou et al., 2018). Cuticle-related genes in *T. castaneum* can have diverse functions in development and physiology (Arakane et al., 2009; Jasrapuria et al., 2012; Zhu et al., 2008), and as noted above, cuticular modifications have been associated with organophosphate and pyrethroid resistance in other insect species (Lilly et al., 2016; Tandonnet et al., 2020; Zimmer et al., 2017). Here, we identified several genes annotated to chitinase, peritrophic matrix, and cuticular proteins differentially expressed with pesticide selection and exposure. In the absence of pesticides, the Coffee population constitutively differentially expressed many cuticle-associated genes and pesticide-selected Coffee larvae also upregulated a number of these. When exposed to pesticides, all three populations differentially expressed cuticle-related genes, and Dorris, in particular, upregulated many. Overall, populations differentially expressed a greater number of chitinases and cuticle associated genes in response to organophosphate selection and exposure compared to pyrethroid interactions. Yet, all organophosphate- and pyrethroid-selected populations commonly differentially expressed cuticle-related genes when exposed to pesticides, raising the possibility that these genes could contribute to multiple-pesticide resistance. Changes in cuticular barriers, including in the gut, are likely to contribute to the increased resistance to pesticides in all pesticide-selected populations examined here.

In addition to physiological modifications, metabolic detoxification may contribute to patterns of cross- or multiple resistance in disease vector and insect pests (Carletto et al., 2010; Carvalho et al., 2013; Edi et al., 2014; Yunta et al., 2019). Oxidative or hydrolytic processes often underlie cross-resistance patterns observed in insects (Basit, 2019; Meinke et al., 2021), and common differential expression of esterases, GSTs, and P450s is often observed between organophosphate and pyrethroid resistant populations (Carletto et al., 2010; Carvalho et al., 2013; Edi et al., 2014; Julio et al., 2017; Yunta et al., 2019). Here, pesticide selected populations varied in the genes commonly differentially expressed with pesticide regime and exposure indicating variation in the possibility for cross-resistance in populations. If cross-resistance is possible for populations, Snavely pesticide-selected populations may rely more on canonical detoxification genes like P450s, GSTs, and UGTs, while Coffee and Dorris may utilize more cuticle-related adaptations. Future research investigating the patterns of cross-resistance in these pesticide resistant populations will help illuminate the mechanisms underlying the evolution of cross-resistance to multiple pesticides.

Insecticide resistance is often associated with fitness costs (Hanai et al., 2018; Homem et al., 2020; Rinkevich et al., 2013; Tieu et al., 2016), but costs are not ubiquitous (ffrench-Constant and Bass, 2017; Freeman et al., 2021) and those associated with metabolic resistance are not as well characterized (Smith et al., 2021). We found that most pyrethroid-selected populations did not demonstrate substantial costs to development. Interestingly, P450-associated pyrethroid resistance in *Anopheles funestus* has been associated with developmental costs (Tchouakui et al., 2020). Here, pyrethroid-resistant populations commonly downregulated P450s without widespread, strong developmental costs. After organophosphate selection, three populations experienced slower development, in agreement with other studies that found developmental costs associated with organophosphate resistance (Abbas et al., 2014; Assogba et al., 2016; Diniz et al., 2015). In comparison to the other two populations chosen for RNA-seq, the Snavely OP-selected population eclosed about a day later (compared to Snavely control) and differentially expressed a greater number of canonical detoxification genes, especially those involved in oxidoreductase activity. However, in the absence of functional genomic experiments, we cannot directly associate specific mechanisms with costs. While we found no fecundity-related costs associated with continued organophosphate selection, the OP-resistant populations exposed to pesticides had higher fecundity relative to unexposed control populations, indicating a context-dependent reproductive advantage. Resistance without associated fitness costs is not uncommon (Freeman et al., 2021) and can contribute to the rapid evolution and spread of beneficial alleles and the maintenance of resistance even in the absence of pesticides (Bird et al., 2020; David et al., 2018; Vila-Aiub et al., 2014). Here, the development of resistance to multiple pesticide types without ubiquitous, strong fitness costs is likely to influence the spread of beneficial adaptations among and between populations.

Across the world, *T. castaneum* populations have developed resistance to all major classes of pesticides used against them (David W Hagstrum and Phillips, 2017), becoming a model for studies on insecticide resistance (Adamski et al., 2019; Zhu et al., 2010). Notably, tandem duplications of cytochrome P450s in the *T. castaneum* genome resulted in expansions of the CYP3 and CYP4 clans known to be involved in environmental and xenobiotic responses and are thought to contribute to *T. castaneum*’s ability to rapidly evolve pesticide resistance (Tribolium Genome Sequencing Consortium, 2008). Here, we identified a large number of P450s, most in the CYP3 clan, differentially expressed in response to pesticide selection and exposure. Interestingly, six of the differentially expressed CYP6BQ genes lie within a single cluster on chromosome LG4 (Zhu et al., 2013). Glutathione S-transferases have also undergone species-specific expansions in *T. castaneum* (Shi et al., 2012), and we identified several GSTs, particularly Delta and Sigma class GSTs, differentially expressed with pesticide selection and exposure. These two classes of GSTs have been implicated in insecticide resistance and oxidative stress responses in *T. castaneum* (Shi et al., 2012; Song et al., 2021, 2020). Our results provide support for the importance of diverse P450s and GSTs in the ability of these beetles to quickly adapt to diverse xenobiotics.

The gene expression patterns exhibited by different populations exposed to the same pesticide selection regime represent both similar (as in Pyr-selected) and different (OP-selected) avenues towards achieving resistance. Though all of our populations were initially reasonably susceptible to both pesticides, we do not know the historical use of pesticides that populations may have encountered in their original field conditions. Adaptations can arise from *de novo* mutations or from standing genetic variation, and it is possible that the transcriptional changes observed in this experiment are reflective of the presence of protective resistance alleles that had been maintained in the population in the absence of pesticide exposure (Bailey and Bataillon, 2016; Long et al., 2015). For example, compared to the other populations, Coffee constitutively differentially expressed many cuticle-related genes that were further differentially expressed after selection to survive pesticide exposure. Future experiments are needed to decipher the genetic architecture and regulatory networks underlying the polygenic adaptations associated with resistance (Barghi et al., 2020; Lu et al., 2020; Signor and Nuzhdin, 2018). Furthermore, mechanisms of evolved pesticide resistance and pesticide exposure are likely to interact or influence immunity against pathogens or other microorganisms (Calhoun et al., 2021; James and Xu, 2012; Minetti et al., 2020). Here, the differential expression of serine endopeptidases and other immune related genes in pesticide-selected populations may point to exciting lines of future research investigating interactions between resistance, pesticide exposure, and immunity against pathogens.

Overall, we found that all populations evolved substantial resistance to pesticides after several generations of selection but achieve this advantage through different putative mechanisms, not all of which are associated with fitness costs. Our results highlight that the degree of common changes in gene expression is dependent on the stressor, even among neurotoxic pesticides. Moreover, the level of parallel changes responding to both pesticides is variable between populations. These findings contribute to our understanding of the diversity of metabolic processes that facilitate adaptation.

## Methods

### T. castaneum Control and Pesticide-selected Colony Maintenance

We used six *T. castaneum* populations (Adairville, Coffee, Dorris, RR, Snavely, WF Ware) collected from the southeast USA (Jent et al., 2019) in experiments to assess how evolved pesticide resistance impacts organismal fitness and molecular interactions between pesticides and pathogens. These populations have been maintained under laboratory control conditions (standard diet of whole wheat flour + 5% yeast; 30°C; 70% humidity; in the dark) for at least two years prior to the start of the experiment.

For each of the six ancestral *T. castaneum* populations, we initially exposed approximately 200 larvae to control conditions or to pesticides (organophosphate [OP] or pyrethroid [Pyr]) (Fig. 1). Initial exposure doses were calculated as one/tenth the manufacturer’s recommended dose (OP [Malathion 50% EC, Southern Ag]- 0.103 mg/ml; Pyr [Demon WP, 40% cypermethrin, Syngenta]- 0.0251 mg/ml). We approximated LC50 pesticide concentrations for ancestral populations using dose response curves (Suppl. Figs. 8A, B), and after the second-generation transfer, we adjusted pesticide concentrations to LC50 concentrations (5.14 mg/ml OP and 0.188 mg/ml Pyr). Pesticide solutions diluted in DI water were combined with 0.15 g/ml standard diet, and approximately 10 ml of the pesticide- or water-diet slurry (for controls) was added to 1 L plastic colony containers and allowed to dry overnight at 55°C. Approximately 200 pupae and larvae were allowed to feed on the pesticide diet for one week and then colonies were supplemented with 100 ml of fresh standard diet. Individuals were transferred to new pesticide exposure containers every four weeks. After six generations of selection, we tested for increased pesticide resistance in F2 larvae from the pesticide-regime populations and found these populations had increased survival against pesticides compared to control populations (Suppl. Figs. 8C, D).

### Effects of Pesticide Selection and Exposure on Survival, Development, and Gene Expression

After eight generations of selection, we prepared same-age F2 larvae from pesticide- and control-regime populations under control conditions to ensure there were no parental effects of pesticide exposure (Fig. 1), and measured the effects of OP and Pyr pesticides on survival and development. Approximately 200 adults from Gen. 8 pesticide- and control-regime populations were allowed to reproduce in 100 ml of control diet for three days. These F0 adults were removed, and offspring were allowed to develop to F1 adults. To generate F2 experimental larvae, we placed 80 F1 adults in 100 mm Petri dishes with control diet and allowed them to reproduce for 24 hours (per population: 5 replicates of F1 control-regime adults, 4 replicates of F1 pesticide-regime adults).

We prepared approximate LC50 pesticide diets as described above. Control diets were prepared by combining 0.15 g/ml standard diet in DI water. We prepared plates to monitor individual larvae by pipetting 50 µl of the control or pesticide diet slurry into each well of 96-well, flat-bottomed cell culture plates. Plates were covered and allowed to dry overnight at 55°C. Twelve-day old larvae (∼4 mm long) were placed on oral diet plates (Fig. 1; n = 96/ population/ treatment). Half of the plate (n = 48) was monitored daily for three weeks for survival and development. Survivors in the remaining half were sampled after two days of exposure and used for RNAseq analyses (described below).

### Effects of Organophosphate Selection and Exposure on Fecundity

To determine whether the evolution of organophosphate resistance is costly to traits associated with fitness, we tested effects of selection regime and pesticide exposure on fecundity. After selection for 16 generations, we produced an F1 generation for control and OP-selected lines from all six populations described above. After one generation of relaxed selection pressure, we collected 160 adults from each of the twelve selection lines (six selected and six control lines) and distributed them to two replicates of 80 adults each. The beetles laid eggs for 24 h in a Petri dish (100 mm diameter) with *ad libitum* standard diet, before being transferred to a new Petri dish for three total egg-laying periods.

When the larvae were two weeks old (14 days post oviposition for the Snavely populations and 13 days post oviposition for all other populations), we transferred them to 96-well plates containing either the control or OP pesticide diet prepared as described above. Pesticide diet contained 10.28 mg/ml OP for the OP-evolved lines and 1.028 mg/ml OP for the control lines to minimize survivorship bias effects across regimes by limiting pesticide induced mortality to about ten percent. To avoid developmental delays and additional deaths, we moved all surviving larvae to new pesticide free diet discs 54 h after initial exposure. We checked larval survival and pupation daily. We recorded pupal weight and sex on the day of pupation and transferred all individuals to empty wells of new 96-well plates. We added dry standard diet to wells containing freshly eclosed adults.

Five days after all adults had reached maturity, we placed single pairs (n≈20 per population and selection * exposure combination) in a small Petri dish (60mm) with standard diet. After 48 h, we transferred the adults to new Petri dishes, while eggs remained in the flour. We conducted three rounds of oviposition in this manner. After three to four weeks post oviposition, we counted live offspring for each pair and oviposition period. For the analysis, offspring counts from the three oviposition periods were combined to obtain one number representing total reproductive output over six days.

### Statistical Methods for Fitness Experiments

We measured mortality and development time to the adult stage for the first 21 days after treatment exposure. We performed survival analyses using the R package ‘survival’ to investigate the impact of treatments on survival and adult development (Therneau and Grambsch 2000; Therneau 2021; R version 4.0.3). There was very little (0-8%) mortality in the control treatments. Thus, to investigate the impact of selection regime on survival against pesticide treatments, we analyzed the effect of selection regime on survival among pesticide treated groups for each population separately (survival ∼ regime). We first created coxph survival models and used the function cox.zph to test adherence to model assumptions. If the coxph assumptions were invalidated, we used the survreg function to create parametric survival models with all possible distributions for individual population models; the best fit model was selected based on the lowest AIC. Similarly, we investigated the main and interactive effects of selection regime and pesticide treatment on adult development using coxph and parametric survival models for each population separately (adult development ∼ regime*treatment), adding a single ‘dummy’ event to treatments that produced no adults to allow model convergence. Pupation and eclosion rates were similar in each group (data not shown), thus for simplicity we present adult development data.

For the fecundity experiment, we first tested whether the maternal and/or paternal pupal weight influenced fecundity in a GLMM including population as a random factor. We continued the analysis only for the mothers, because we only observed a significant correlation between pupal weight and fecundity for the females. We started with the most complex mixed model including the three-way interaction between pupal weight, OP selection regime and OP exposure (fecundity ∼ weight*regime*exposure + (1|population)). Stepwise model simplification gave us the minimal model including the interaction terms for weight and pesticide exposure as well as pesticide exposure and selection regime (fecundity ∼ weight + regime*exposure +(1|population)). Data were not normally distributed and were overdispersed, so we implemented negative binomial GLM analyses using the R package ‘MASS’ (Venables and Ripley 2002).

### RNA Extractions and Sequencing

To measure changes in gene expression in representative populations, we selected Coffee, Dorris, and Snavely for RNAseq analyses, taking advantage of the well-annotated, published genome for *T. castaneum* (Herndon et al., 2020). For each population, samples for 3’-TagSeq (Lohman et al., 2016) were collected after two days of exposure to oral pesticide diets, prior to the main onset of mortality. For each population-by-treatment group, nine surviving larvae were collected and immediately stored at −80°C until processing. For each group, we pooled three larvae for RNA extraction to generate three biological replicates. Total RNA was extracted using a Qiagen RNeasy Mini kit according to manufacturer protocols. Remaining genomic DNA was removed using the Invitrogen RNaqueous Micro DNase treatment according to manufacturer protocols. Nucleic acid concentration and quality was confirmed using a Nanodrop. We also confirmed nucleic acid concentration and quality of twelve randomly selected DNase-treated RNA samples using a Bioanalyzer (RIN > 9). Single indexed 3’ TagSeq libraries were generated from 1 µg total RNA using the Lexogen QuantSeq 3’ mRNA-Seq Library Prep Kit according to manufacturer’s protocols. Libraries were amplified using 15 PCR cycles. Library concentration and quality were confirmed prior to SE75 sequencing across four runs on the NextSeq platform (VANTAGE, Vanderbilt University).

### RNAseq Analyses

Raw mRNAseq reads were trimmed for adapters and low-quality reads were removed using fastqc (Andrews 2010); a total of 6.8-16.3 million reads per sample were retained (Suppl. Table 5; average reads per sample = 10.7 million). Sequence reads are deposited in the NCBI SRA under the BioProject Accession: PRJNA753142). Trimmed reads were aligned to the *T. castaneum* genome (Tcas5.2) using BWA-MEM (Suppl. Table 5; average reads mapped = 84.98%, s.d. = 0.025; (Li, 2013)) and gene count files for each library were generated using samtools (Li et al., 2009). We used DESeq2 (Love et al., 2014) to perform differential expression analyses and identify significantly differentially expressed genes (padj < 0.05). First, we subset control-regime, control treatment libraries and examined constitutive gene expression differences between populations (count matrix ∼ population). To visualize overall differences between populations, principal component analysis (PCA) plots were generated using the model: count matrix ∼ population + regime*treatment. Last, we analyzed each population was analyzed separately to identify significantly differentially expressed genes (padj < 0.05) with respect to the main and interactive effects of selection regime and pesticide treatment (count matrix ∼ regime*treatment.) Differentially expressed transcripts were annotated using the Ensembl *T. castaneum* database and the R package ‘biomaRt’ (Durinck et al., 2009). Gene ontology (GO) enrichment analyses for differentially expressed gene sets were performed using the online Gene Ontology Resource (Ashburner et al., 2000; Gene Ontology Consortium, 2021). We manually curated the annotated gene lists to categorize differentially expressed transcripts into groups potentially impacted by pesticide exposure and resistance (Suppl. Table 6).

## Supplemental Figures

**Suppl. Figure 1.**
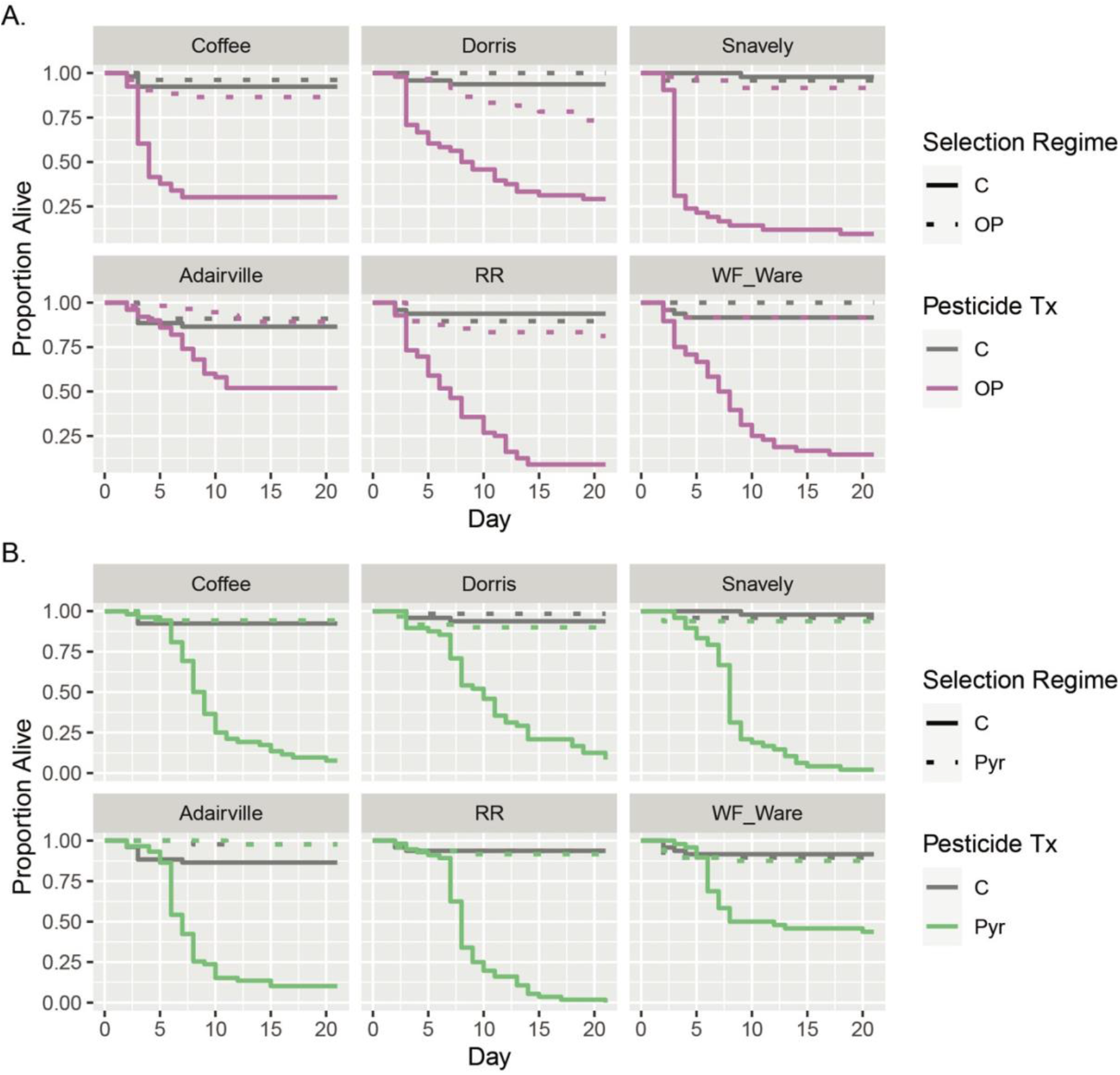
Effects of pesticide selection and exposure on survival in six *T. castaneum* populations. **A.** Effects of organophosphate (OP) selection and exposure on daily proportion of surviving individuals. **B.** Effects of pyrethroid (Pyr) selection and exposure on daily proportion of surviving individuals.

**Suppl. Figure 2.**
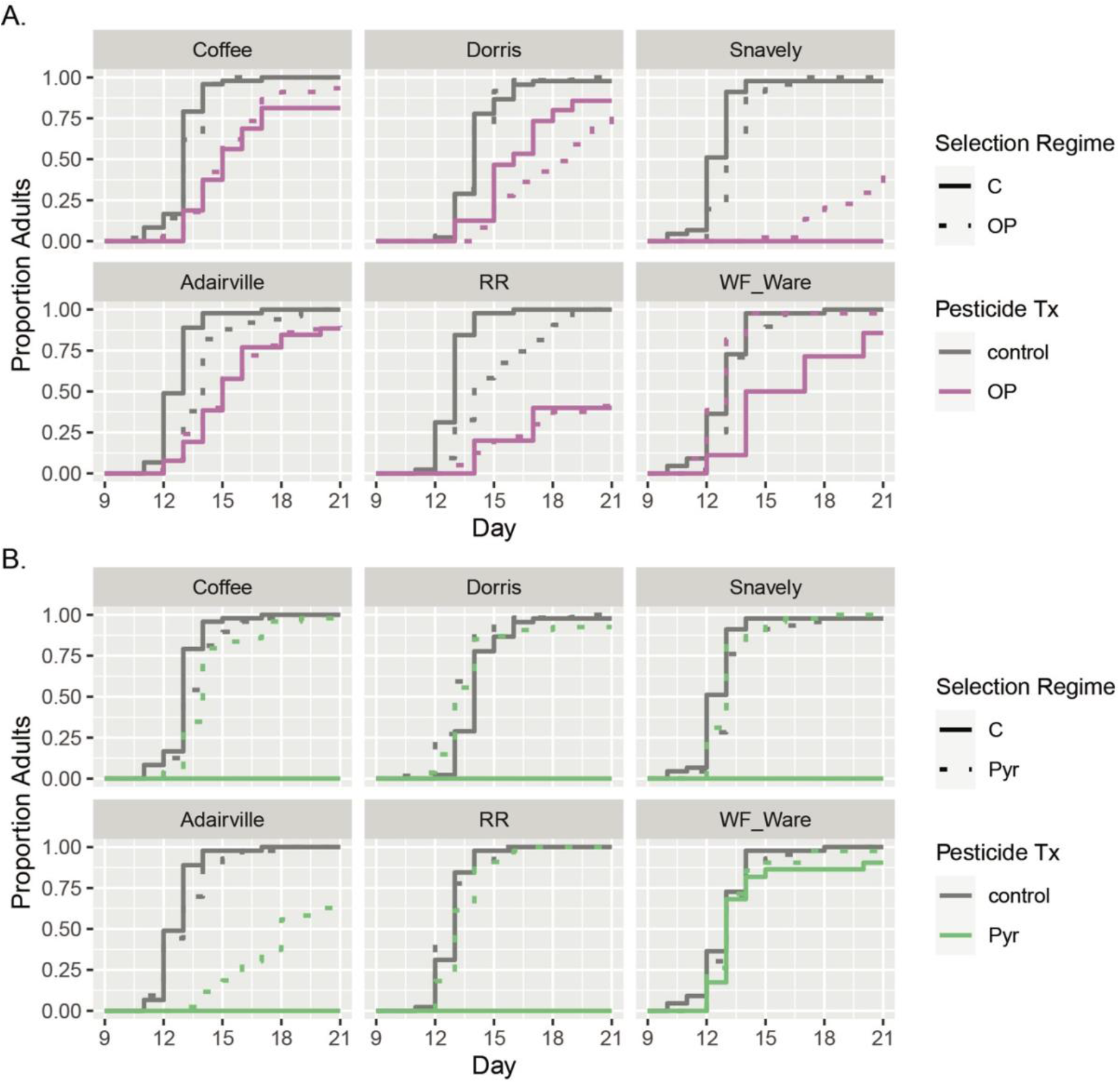
Effects of pesticide selection and exposure on adult development in six *T. castaneum* populations. **A.** Effects of organophosphate (OP) selection and exposure on proportion of eclosed adults. **B.** Effects of pyrethroid (Pyr) selection and exposure on daily proportion of eclosed adults.

**Suppl. Figure 3.**
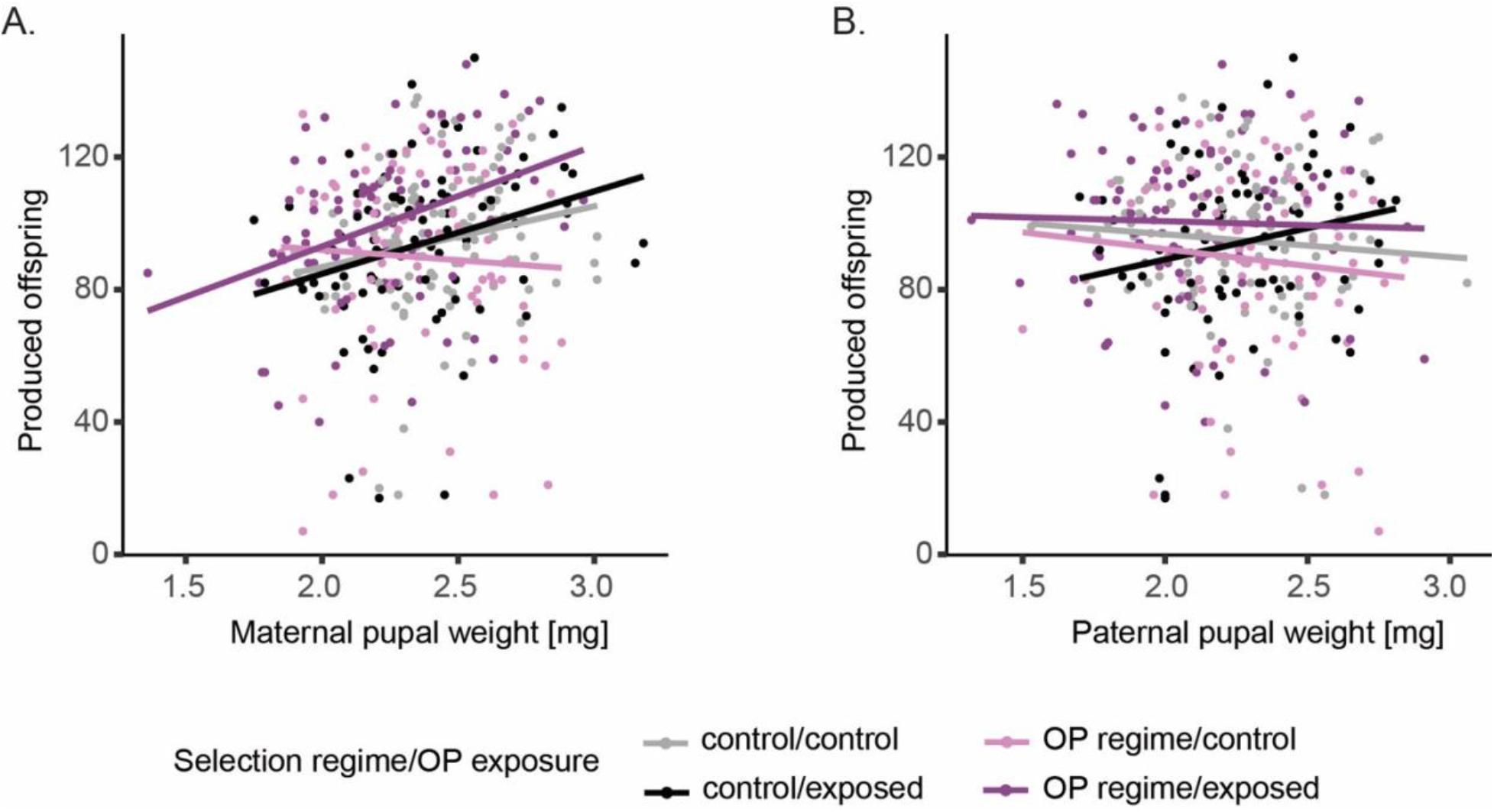
Effects of OP selection regime and exposure, and **A.** female and **B.** male pupal weight on fecundity (produced live offspring). Dots represent offspring per individual mating pair.

**Suppl. Figure 4.**
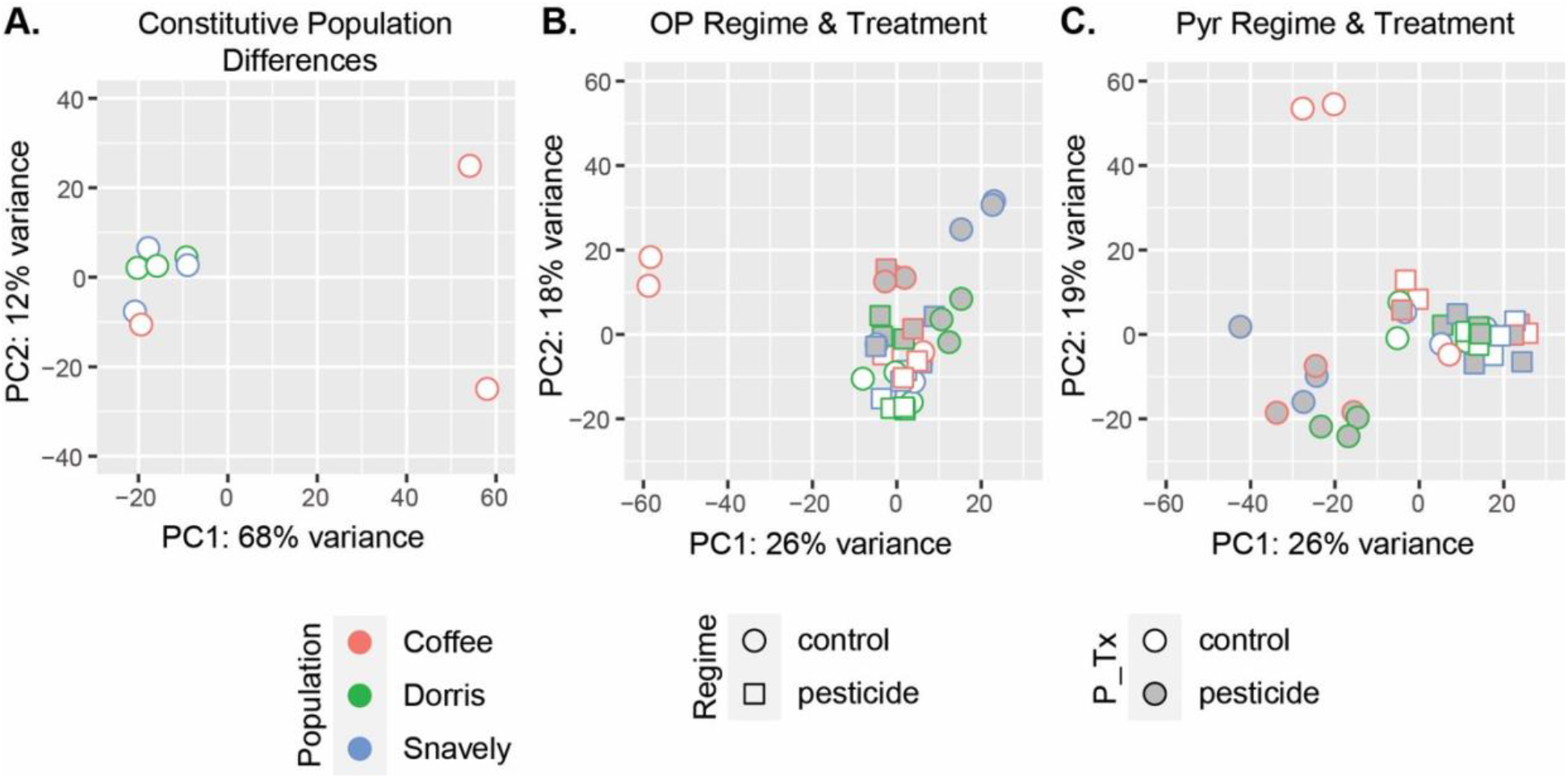
Principle components analysis (PCA) plots. **A.** Constitutive population differences, **B.** control- and OP-regime populations exposed to control and OP treatments, and **C.** control- and Pyr-regime populations exposed to control and Pyr treatments.

**Suppl. Figure 5.**
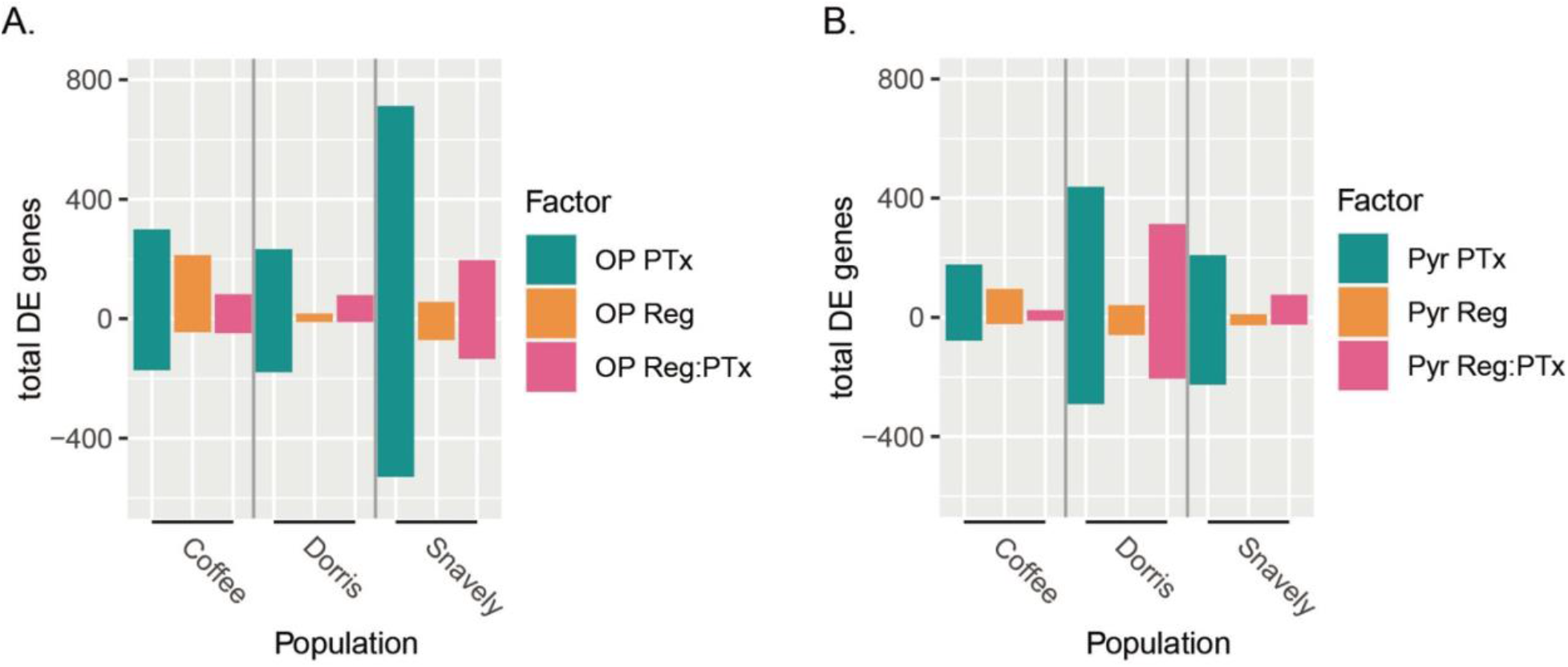
Total number of genes significantly differentially expressed (padj < 0.05 after FDR correction) with respect to the main and interactive effects between pesticide selection and exposure. DE gene counts between control and pesticide treatments (PTx), control and pesticide selection regimes (Reg), and those responding differently in selected populations compared to control populations when exposed to pesticides (Reg:PTx) are show for each population (upregulated are positive counts, downregulated are negative counts). **A.** Effects of organophosphate (OP) selection and exposure. **B.** Effects of pyrethroid (Pyr) selection and exposure.

**Suppl. Figure 6.**
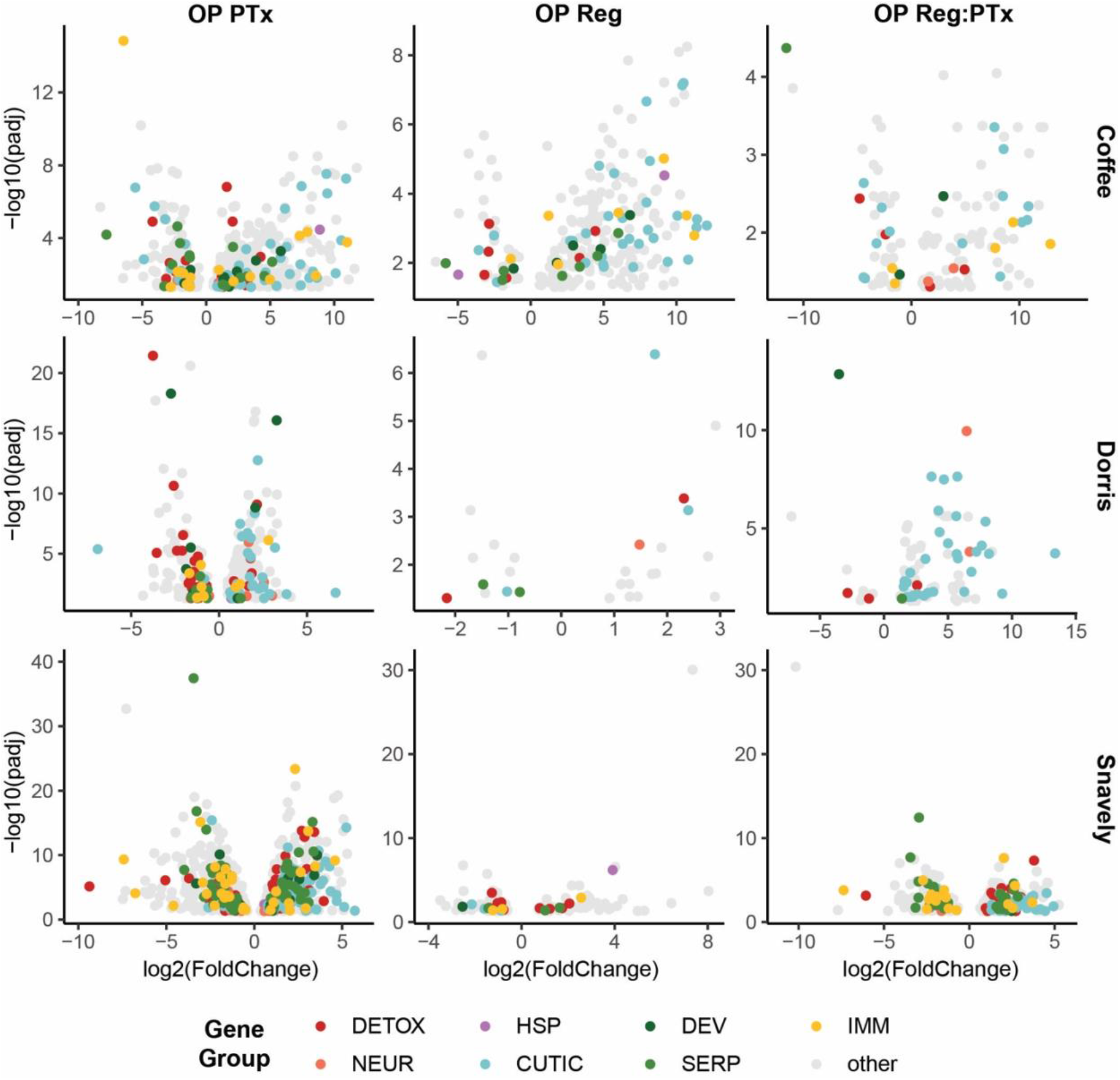
Volcano plots of all genes that are significantly differentially expressed with organophosphate selection and exposure (padj < 0.05 after FDR correction). OP PTx: genes differentially expressed with OP exposure compared to control treatment; OP Reg: genes differentially expressed in OP selection regime compared to control regime; OP Reg:PTx: genes differentially expressed with OP exposure in OP regime compared to control regime. DETOX = canonical detoxification, NEUR = neural function, HSP = heat shock protein, CUTIC = cuticle-related genes, DEV = development genes, SERP = serine protease/ serpin, IMM = immunity related genes, other = all other transcripts.

**Suppl. Figure 7.**
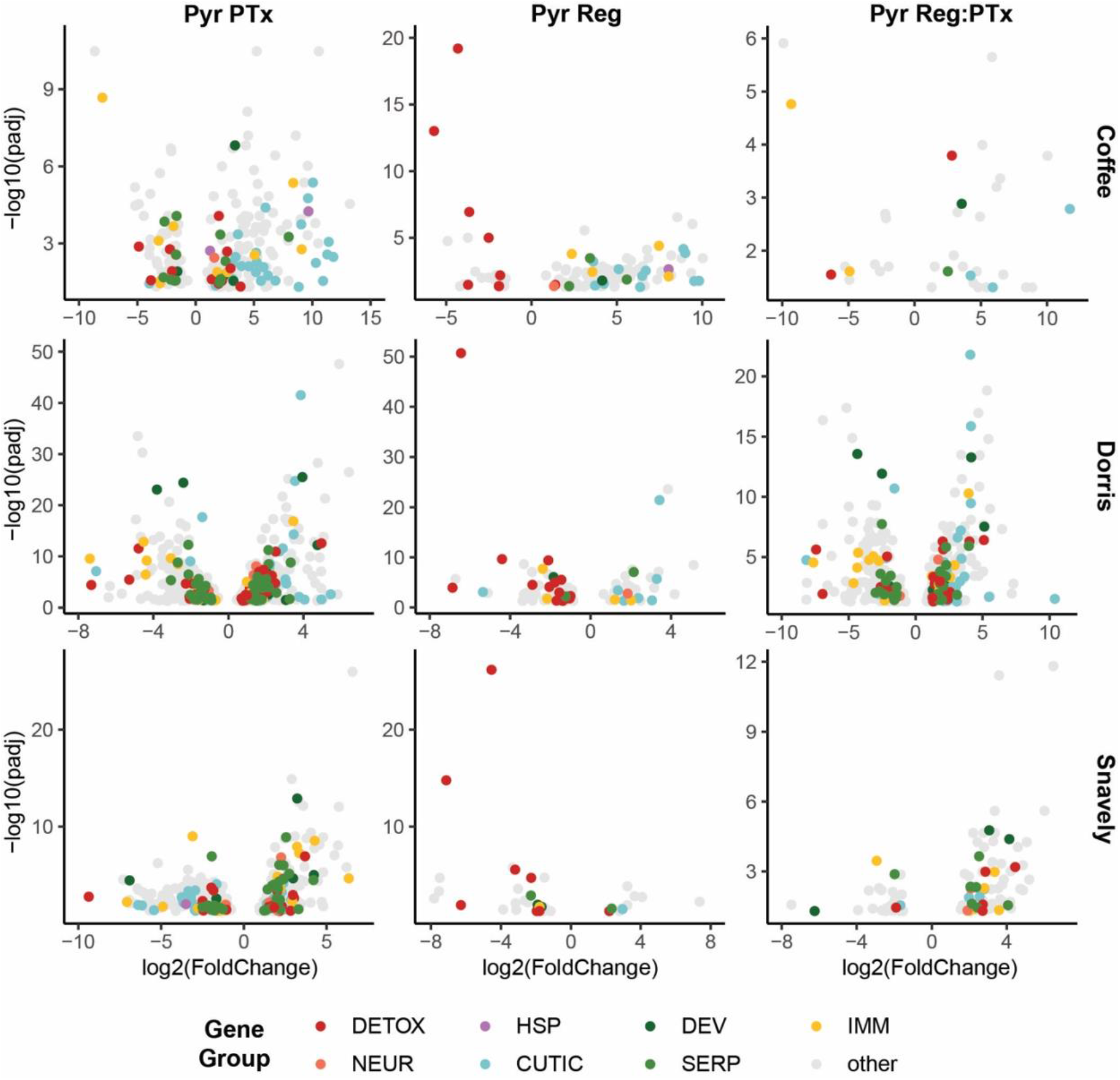
Volcano plots of all genes that are significantly differentially expressed with pyrethroid selection and exposure (padj < 0.05 after FDR correction). Pyr PTx: genes differentially expressed with Pyr exposure compared to control treatment; Pyr Reg: genes differentially expressed in Pyr selection regime compared to control regime; Pyr Reg:PTx: genes differentially expressed with Pyr exposure in Pyr regime compared to control regime. DETOX = canonical detoxification, NEUR = neural function, HSP = heat shock protein, CUTIC = cuticle-related genes, DEV = development genes, SERP = serine protease/ serpin, IMM = immunity related genes, other = all other transcripts.

**Suppl. Figure 8.**
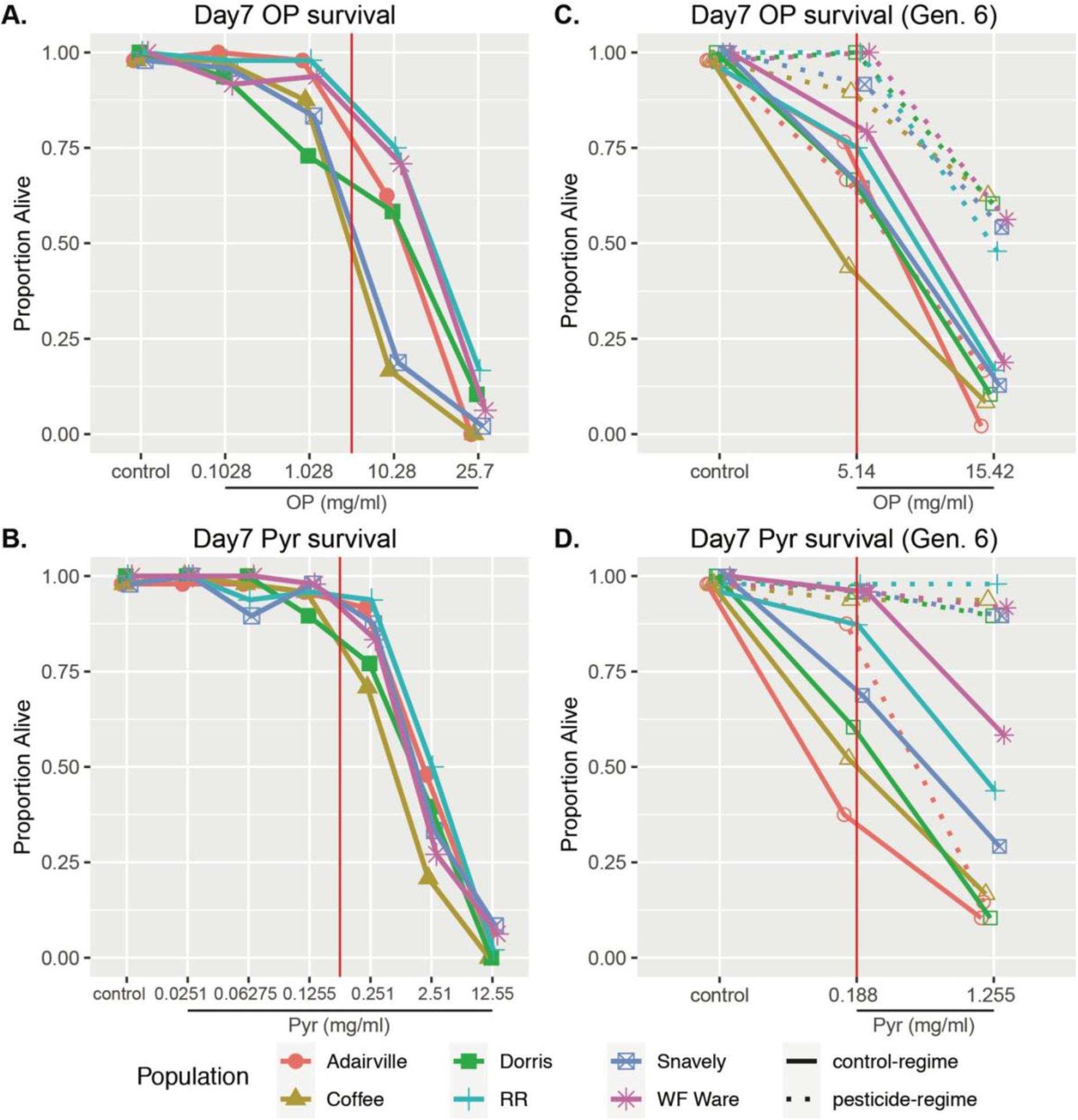
Preliminary dose response curves in ancestral, control-regime populations and testing of evolved resistance in pesticide selected populations. Survival after seven days of treatment exposure is shown. **A.** Organophosphate (OP) survival dose response curve for control-regime populations. **B.** Pyrethroid (Pyr) survival dose response curve for control-regime populations. **C.** Testing for increased survival in Generation 6 organophosphate-regime populations compared to control-regime populations. **D.** Testing for increased survival in Generation 6 pyrethroid-regime populations compared to control-regime populations. Approximate LC50 OP and Pyr doses used in experiments are shown with red lines.

## Supplemental Tables

**Suppl. Table 1.** Effect of selection regime on survival among pesticide treated groups (base level is control treatment).

**Suppl. Table 2.** Main and interaction effects of pesticide exposure and selection on adult development (compared to control regime and treatment). PTx = pesticide treatment (base level is control), Reg = evolution regime (base level is control regime), Reg:PTx = the interaction of evolution regime and pesticide treatment.

**Suppl. Table 3.** Effects of male and female pupal weight on fecundity regardless of OP selection regime and OP exposure treatment groups. GLMM Fecundity ∼ Male weight + Female weight + (1| Population)

**Suppl. Table 4.** Effects of OP selection regime and exposure, and female pupal weight on fecundity (produced offspring). GLMM: Fecundity ∼ Regime*Tx + Female weight + (1| Population)

**Suppl. Table 5.** RNAseq sample details.

**Suppl. Table 6.** Full list of differentially expressed transcripts. For each factor, the log_2_ fold change (log2FoldChange) and adjusted p-values (padj) are given. Sample abbreviations are as follows: constitutively differentially expressed between populations: Co_Do = Coffee vs. Dorris, Co_Sn = Coffee vs. Snavely, Do_Sn = Dorris vs. Snavely; differentially expressed transcripts for organophosphate samples (Co_ = Coffee, Do_ = Dorris, Sn_ = Snavely): OP_PTx = control vs. OP treatment, OP_Reg = control- vs. OP-regime, OP_RegPTx = regime and pesticide treatment interaction; differentially expressed transcripts for pyrethroid samples (Co_ = Coffee, Do_ = Dorris, Sn_ = Snavely): Pyr_PTx = control vs. Pyr treatment, Pyr_Reg = control- vs. Pyr-regime, Pyr_RegPTx = regime and pesticide treatment interaction.

**Suppl. Table 7.** Full list of GO enrichment results for organophosphate DE lists. DE factors: OP_PTx = control vs. OP treatment, OP_Reg = control- vs. OP-regime, OP_Reg:PTx = OP- regime and OP treatment interaction. GO function: bio = biological process, mol = molecular function, cell = cellular component.

**Suppl. Table 8.** Full list of GO enrichment results for pyrethroid DE lists. DE factors: Pyr_PTx = control vs. Pyr treatment, Pyr _Reg = control- vs. Pyr -regime, Pyr _Reg:PTx = Pyr -regime and Pyr treatment interaction. GO function: bio = biological process, mol = molecular function, cell = cellular component.

## Acknowledgements

We would like to thank William Galardi for assistance in preliminary dose response experiments and Siqin Liu for assistance with the fecundity experiment and population maintenance. We are grateful to Juan Luis Jurat-Fuentes for comments on the manuscript. This work is funded by USDA NIFA 2019-67012-29659 provided to S.S.L.B. N.K.E.S. and A.T.T. were supported in part by Alfred P. Sloan Foundation Fellowship FG-2020-12949 to A.T.T.

## Author Contributions

S.S.L.B. and A.T.T. designed the study. S.S.L.B. executed the main fitness and RNAseq experiments and associated statistical analyses. D.S.G. and N.K.E.S. performed the fecundity experiment, and N.K.E.S. executed the fecundity statistical analyses. S.S.L.B., A.T.T., and N.K.E.S wrote the manuscript.

